# *OPA1* and disease-causing mutants perturb mitochondrial nucleoid distribution

**DOI:** 10.1101/2024.02.01.578418

**Authors:** J. Macuada, I. Molina-Riquelme, G. Vidal, N. Pérez-Bravo, C. Vásquez-Trincado, G. Aedo, D. Lagos, R. Horvath, T.J. Rudge, B. Cartes-Saavedra, V. Eisner

**Affiliations:** Facultad de Ciencias Biológicas, Pontificia Universidad Católica de Chile, Santiago, Chile; Institute for Biological and Medical Engineering, Schools of Engineering, Biology, and Medicine, Pontificia Universidad Católica de Chile, Santiago, Chile; Interdisciplinary Computing and Complex Biosystems (ICOS) research group, School of Computing, Newcastle University, Newcastle upon Tyne, UK; Department of Clinical Neurosciences, John Van Geest Centre for Brain Repair, University of Cambridge, Ed Adrian Building, Robinson Way, Cambridge, CB2 0PY, UK; Department of Chemical and Bioprocess Engineering, School of Engineering, Pontificia Universidad Católica de Chile, Santiago, Chile; MitoCare Center for Mitochondrial Imaging Research and Diagnostics, Department of Pathology and Genomic Medicine, Thomas Jefferson University, Philadelphia, PA, USA

## Abstract

Optic atrophy protein 1 (OPA1) mediates inner mitochondrial membrane (IMM) fusion and cristae organization. Mutations in OPA1 cause autosomal dominant optic atrophy (ADOA), a leading cause of blindness. Cells from ADOA patients show impaired mitochondrial fusion, cristae structure, bioenergetic function, and mitochondrial DNA (mtDNA) integrity. The mtDNA encodes electron transport chain subunits and is packaged into nucleoids spread within the mitochondrial population. Nucleoids interact with the IMM, and their distribution is tightly linked to mitochondrial fusion and cristae shaping. Yet, little is known about the physio-pathological relevance of nucleoid distribution. We studied the effect of OPA1 and ADOA-associated mutants on nucleoid distribution using high-resolution confocal microscopy. We applied a novel model incorporating the mitochondrial context, separating nucleoid distribution into the array in the mitochondrial population and intramitochondrial longitudinal distribution. Opa1-null cells showed decreased mtDNA levels and nucleoid abundance. Also, loss of Opa1 lead to an altered distribution of nucleoids in the mitochondrial population, loss of cristae periodicity, and altered nucleoids to cristae proximity partly rescued by OPA1 isoform 1. Overexpression of WT OPA1 or ADOA-causing mutants c.870+5G>A or c.2713C>T in WT cells, showed perturbed nucleoid array in the mitochondria population associated with cristae disorganization. Opa1-null and cells overexpressing ADOA mutants accumulated mitochondria without nucleoids. Interestingly, intramitochondrial nucleoid distribution was only altered in Opa1-null cells. Altogether, our results highlight the relevance of OPA1 in nucleoid distribution in the mitochondrial landscape and at a single-organelle level and shed light on new components of ADOA etiology.

## INTRODUCTION

Mitochondria are critical to intracellular bioenergetic balance (1) and their genome, the mtDNA, encodes genes required to maintain the function of the electron transport chain (ETC). MtDNA maintenance defects are a hallmark of bioenergetic dysfunction in several diseases and aged tissue (2,3). In humans and mice, mtDNA is packaged into 70-110 nm ellipsoid DNA-protein complexes named nucleoids (4,5) through interaction with its main protein component, the Mitochondrial Transcription Factor A (TFAM) (6,7).

The maintenance and distribution of mtDNA are critical for cellular physiology (8–13) and are impaired during disease (14–19). Mitochondria undergo constant fusion/fission cycles, which allow the exchange of its components and supports mtDNA maintenance (1,8,20). In mammalian cells, nucleoids are semi-regularly distributed inside the mitochondrial population (5,21,22) and, in yeast, it was proven that regular distribution is necessary to accurately control nucleoid segregation to daughter cells (23). Furthermore, nucleoid positioning on the sites of mitochondrial fission can ensure mtDNA inheritance to daughter mitochondria (24). Regardless of the relevance of mtDNA abundance and maintenance, little is known about the physio-pathological relevance of nucleoid distribution in the mammalian mitochondrial population, partly because of the challenge of studying nanoscopic structures (25).

Nucleoids physically interact with the inner mitochondrial membrane (IMM), and are interspersed between the IMM cristae (4,26,27). Cristae structure is tightly regulated by multiple proteins, such as the ATP synthase (28), the Mitochondrial Contact Site and Cristae Organization System (MICOS) (29,30), and OPA1 (31–33). OPA1 is a GTPase also involved in various processes, including mitochondrial fusion and fission (34–36). Furthermore, the lack of Mgm1, the OPA1 yeast ortholog, leads to mtDNA depletion (37,38) and *Opa1*^*-/-*^ cells show a decrease in mtDNA and nucleoid number (39–41), and display mitochondria without nucleoids (39). Finally, OPA1exon4b isoforms 3, 5, 6 and 8 physically interact with mtDNA to regulate its maintenance (39,42). Yet, the role of OPA1 in intramitochondrial nucleoid distribution in mammalian cells under pathophysiological conditions is under explored.

*OPA1* mutations lead to Autosomal Dominant Optic Atrophy (ADOA, MIM 165500), a disease that causes progressive loss of retinal ganglion cells resulting in optic nerve degeneration and blindness (43). About 20% of ADOA patients manifest a more severe syndromic form of the disease, classified as ADOA plus (ADOA+, MIM: 125250) (44,45). Patients bearing mutations at the GTPase domain of OPA1 are more likely to develop ADOA+ (45). Skeletal muscle samples from ADOA patients display defects in mtDNA integrity and distribution with bioenergetic dysfunction markers, as well as IMM ultrastructure alterations (18,46,47). Accordingly, our previous studies showed that OPA1 ADOA-causing mutants in the GTPase or GTPase effector domain (GED) display distinct effects on IMM ultrastructure and mitochondrial fusion and fission dynamics (48). Nevertheless, it is unclear if these changes have an impact on mtDNA distribution and maintenance.

In this work we studied the effect of OPA1 and different ADOA-causing mutants on nucleoid distribution. We used *Opa1*^*-/-*^ MEF and overexpression of OPA1 mutants on WT MEF to study nucleoid distribution and nucleoid-cristae proximity. Our data showed that the lack of OPA1 and the overexpression of OPA1 or ADOA-causing mutants perturbed nucleoid cluster distribution within the mitochondrial population and disrupted cristae morphology. Furthermore, nucleoid-cristae proximity is perturbed in *Opa1*^*-/-*^ cells which might contribute to local OXPHOS dysfunction. This work introduces new clues on the influence of OPA1 over mitochondrial nucleoid distribution and provides novel insights on the etiology of the mtDNA alterations as well as the local bioenergetic dysfunction found in ADOA patients.

## RESULTS

### Opa1 determines nucleoid abundance and distribution

We studied the role of OPA1 in nucleoid abundance using *Opa1*^*-/-*^ MEFs and rescue through expression of human OPA1 isoform 1. First, we quantified the mtDNA copy number and found a ∼70% decrease in mtDNA levels in *Opa1*^*-/-*^ cells, which was partially rescued upon acute expression of OPA1, and completely rescued in our *Opa1*^*-/-*^ + OPA1 stable cell line (+ p-lenti OPA1 WT) (**Fig. 1A**). To accurately evaluate nucleoid distribution, we propose a novel model where we measure: 1) the nucleoids array in the mitochondrial population, which describes the density and spread of nucleoids within the mitochondrial network and includes the mitochondria with and without nucleoids and, 2) intramitochondrial longitudinal distribution, which describes nucleoid localization in relation to the length of single organelles (**Fig. 1B**). To apply our model, we developed a manual approach and a novel unbiased semi-automated method called MiNuD (Mitochondrial Nucleoids Distribution), as described in Methods (**Fig. S1)**.

**Figure 1.**
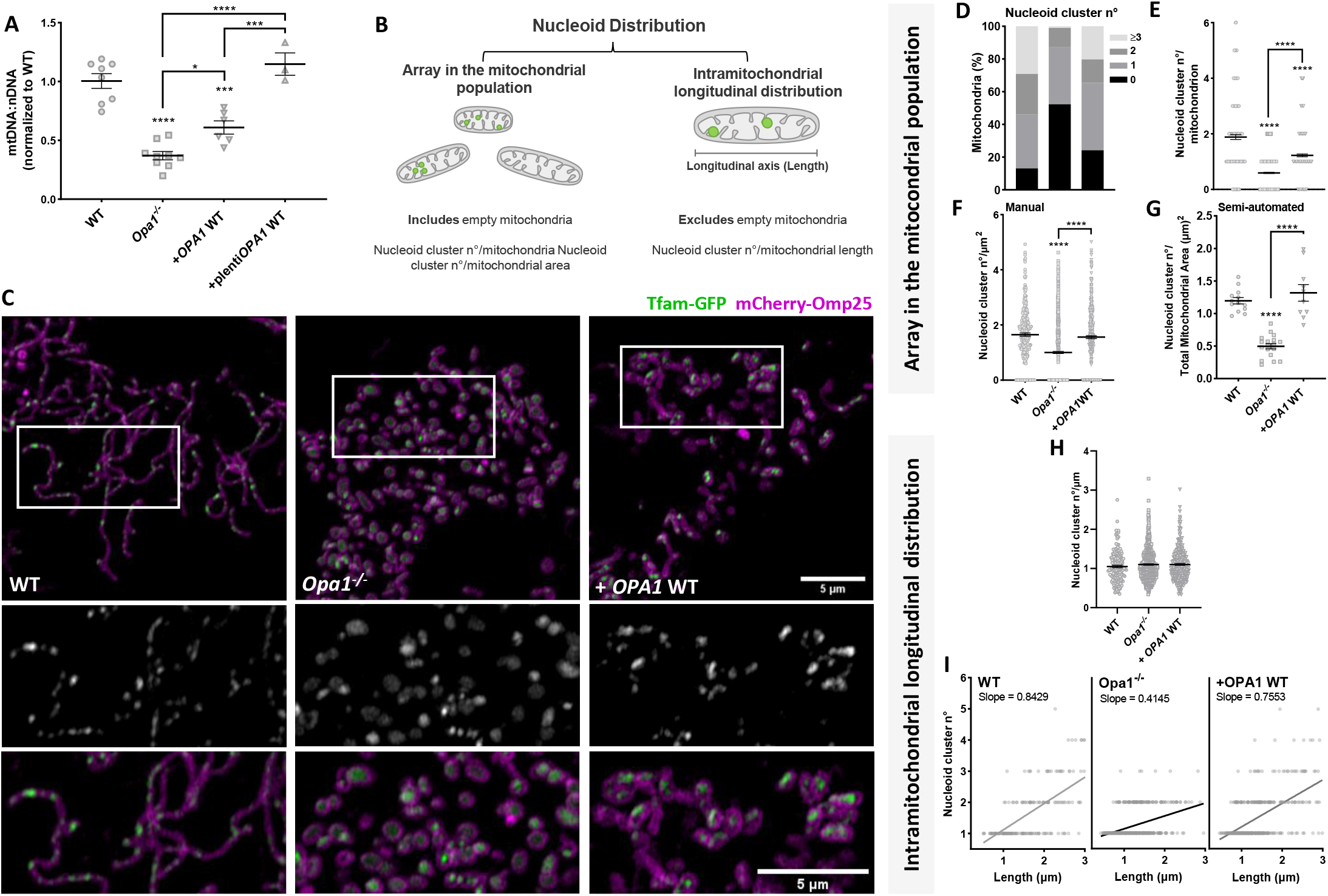
Opa1 determines nucleoid abundance and array in the mitochondrial population and intramitochondrial longitudinal distribution. **(A)** Mitochondrial DNA abundance in WT MEF, *Opa1*^*-/-*^ and *Opa1*^*-/-*^ cells stably expressing a lentiviral plasmid carrying OPA1 cDNA, quantified by qPCR and expressed as mt-Nd4 to Gapdh ratio. Data are mean ± SEM of ≥ 3 independent experiments. Scale bar = 1μm. **(B)** Schematic representation of the components of nucleoid distribution. Scale bar = 500 nm. **(C)** Representative images of WT MEF, *Opa1*^*-/-*^ and *Opa1*^*-/-*^ cells with WT OPA1 acute expression, co-transfected with mCherry-Omp25 (OMM) and Tfam-GFP (nucleoids) cDNA. Bottom panel, insets of Tfam-GFP (top) and merge (bottom). **(D)** Percentage of mitochondria with different number of nucleoid clusters. **(E)** Mean nucleoid cluster number per mitochondrion. **(F)** Mean nucleoid cluster number per mitochondrion area, manually quantified. **(G)** Number of nucleoid clusters per total mitochondrial area, semi-automatically quantified by the MiNuD algorithm. For *Opa1*^*-/-*^ cells, DNA was labeled using PicoGreen, and mitochondria were labeled with mCherry-Omp25 (OMM) or MitoTracker Deep Red. **(H)** Mean nucleoid cluster number per mitochondrion length. **(I)** Simple regression of the relationship between nucleoid cluster number and mitochondrion length in individual organelles. Data are mean ± SEM of ≥ 281 objects from ≥ 15 cells of ≥ 3 independent experiments, and for G data are mean ± SEM of ≥ 10 cells of ≥ 2 independent experiments (^****^ p<0.0001; ^***^ p<0.0005; ^*^ p<0.05)

We applied our semi-automatic approach to quantify nucleoid distribution in WT MEF using two strategies to label nucleoids: i) direct dsDNA labeling with PicoGreen or ii) Tfam-GFP (49). We performed live-cell imaging at a high-resolution confocal microscope, with sufficient resolution to identify nucleoid clusters, rather than individual objects (50). Both techniques showed “point-like” signals with apparent uniform distribution throughout the mitochondrial network (**Fig. S2A**), and comparable number of nucleoid clusters per total mitochondrial area (**Fig. S2B**). Considering that PicoGreen displayed a nuclear stain (**Fig. S2A**), we continued our study with Tfam-GFP, excluding cells that show signs of major overexpression such as nuclear Tfam-GFP localization (49).

We evaluated mitochondrial nucleoid abundance and distribution using the manual quantification approach described in **Methods** (**Fig. S1**). We studied the number of nucleoids per mitochondria in maximum projections (∼ 1μm thickness) and found that in WT MEF, 13% of mitochondria lack apparent Tfam-GFP foci, thus we classified them as “empty”. We corroborated this result with anti-dsDNA and Picogreen staining (**Fig. S2C**).

*Opa1*^*-/-*^ cells displayed an increase in “empty” organelles (52% vs 13% WT) and a diminished number of nucleoid clusters per mitochondria, consistent with the loss in mtDNA levels (**Fig. 1C-D**). Acute rescue of OPA1 partially recovered the number of “empty” mitochondria (24%) and the number of nucleoid clusters per mitochondria (**Fig. 1C-E**). In addition, we measured the number of nucleoid clusters per mitochondrial area using our manual approach and MiNuD and found the same tendency (**Fig. 1F-G**). Notably, in *Opa1*^*-/-*^ cells, Tfam-GFP showed a combination of bright points and diffuse signals (**Fig. 1C**). The acute rescue of OPA1 re-established the nucleoid cluster phenotype and labeling. Alternative nucleoid labeling with PicoGreen or anti-dsDNA confirmed the low number of nucleoid clusters observed in the *Opa1*^*-/-*^ cells (**Fig. S3A-C**).

Considering the high number of mitochondria lacking apparent Tfam-GFP foci and the changes in morphology shown in *Opa1*^*-/-*^ cells, we evaluated intramitochondrial longitudinal distribution in organelles that: 1) showed Tfam-GFP foci; 2) were shorter than 3 μm, the longest organelles found in *Opa1*^*-/-*^ cells (**Fig. S4**). Interestingly, the average intramitochondrial distribution of one nucleoid cluster per micrometer remained unchanged in *Opa1*^*-/-*^ cells (**Fig. 1H**). To further examine this, we applied a simple linear regression to the number of nucleoid clusters found in each mitochondrion and its length. WT MEF showed a correlation between mitochondria length and nucleoid cluster number that is reduced in *Opa1*^*-/-*^ cells and rescued after acute OPA1 expression (**Fig. 1I**).

Thus, OPA1 acute rescue in *Opa1*^*-/-*^ cells partially recovered mtDNA abundance and the nucleoid cluster array in the overall mitochondrial population. Notably, the absence of OPA1 perturbed the intramitochondrial longitudinal distribution of the nucleoid clusters.

### OPA1 rescues mitochondrial cristae periodicity and nucleoid-cristae proximity in *Opa1*^*-/-*^ cells

As OPA1 is key for IMM fusion and cristae organization (32,40) and the cristae morphology is associated with nucleoid distribution and abundance (10,26,41), we performed transmission electron microscopy (TEM) (**Fig. 2A**).

**Figure 2.**
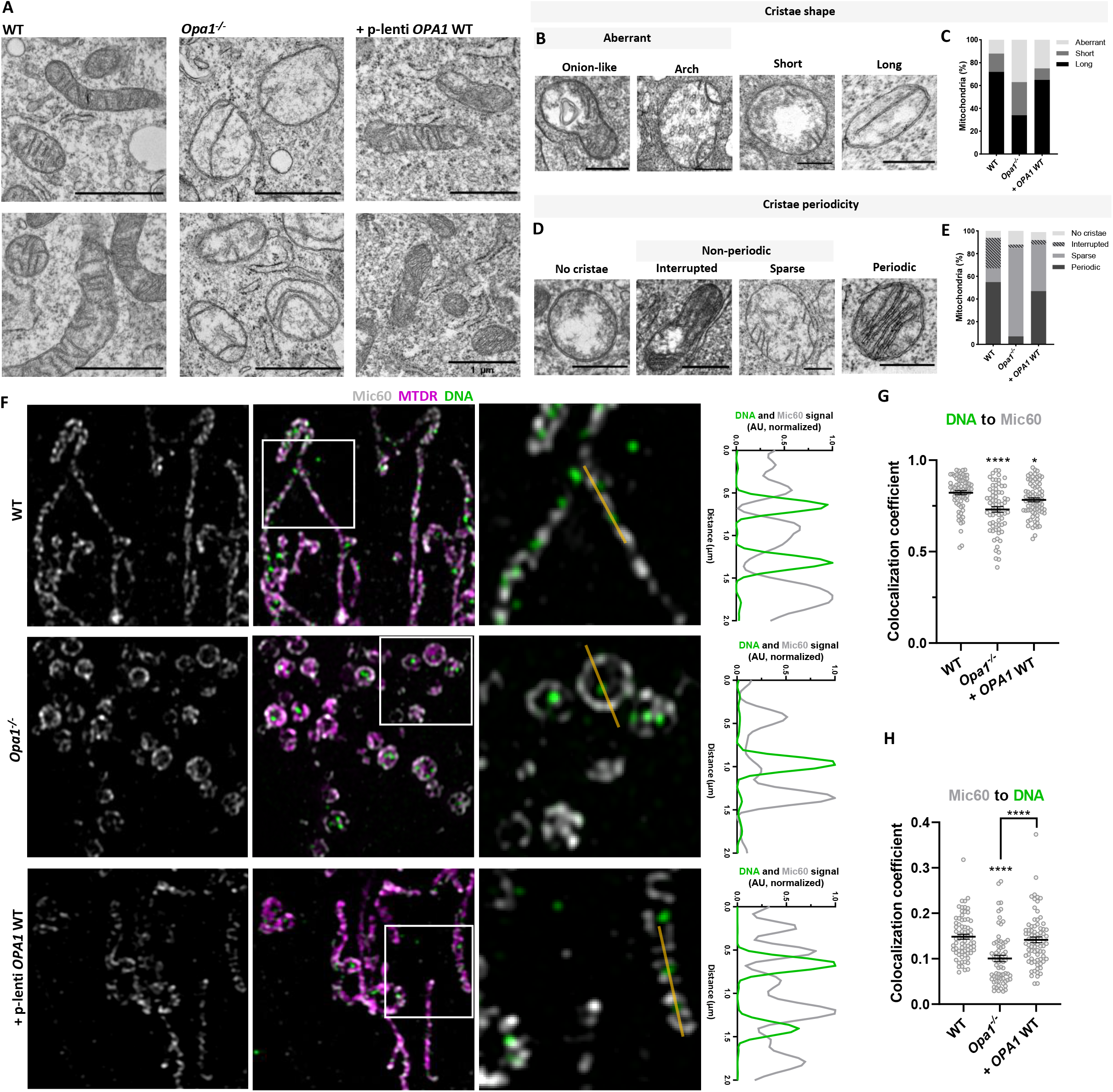
The stable expression of WT OPA1 rescues mitochondrial cristae periodicity and nucleoid-cristae proximity in *Opa1*^*-/-*^ cells. **(A)** Representative TEM images of WT MEF, *Opa1*^*-/-*^ and *Opa1*^*-/-*^ cells stably expressing a lentiviral plasmid carrying OPA1 cDNA. **(B)** Mitochondrial cristae shape classification and **(C)** their quantification. **(D)** Mitochondrial cristae periodicity classification and **(E)** their quantification. Data are from ≥ 98 objects from ≥ 2 independent experiments. **(F)** Representative immunofluorescence images of WT MEF, *Opa1*^*-/-*^ and *Opa1*^*-/-*^ cells stably expressing a lentiviral plasmid carrying OPA1 cDNA. Cells were labeled with MitoTracker Deep Red and antibodies against dsDNA and Mic60. (Right) Intensity profile of dsDNA and Mic60 signals along the yellow line (2 μm). **(G-H)** Colocalization coefficients for dsDNA and Mic60 in the cells shown in F. Each point represents one cell. Data are mean ± SEM from ≥ 70 cells from 4 independent experiments.

Based on the diversity of cristae phenotypes, we classified each mitochondrion according to their cristae shape and periodicity (**Fig. 2B-D**). Mitochondria with no discernible cristae were classified as organelles with “no cristae”. In *Opa1*^*-/-*^ cells, we observed an increase in mitochondria with short (29% vs. 16% WT) and aberrant cristae (37% vs. 12% WT) (**Fig. 2A-C**). The aberrant cristae in *Opa1*^*-/-*^ cells consisted mainly of onion-like and arch phenotypes (**Fig. S3D**). Importantly, the stable expression of OPA1 rescued the abundance of long cristae (65% vs. 34% *Opa1*^*-/-*^) (**Fig. 2A-C**). Similarly, cristae periodicity was disturbed in *Opa1*^*-/-*^ cells. Mitochondria with no cristae were increased (12% vs. 6% WT) as well as non-periodic sparse cristae (78% vs. 12% WT) (**Fig. 2E**). The stable expression of OPA1 partially rescued the abundance of periodic cristae (47% vs. 7% *Opa1*^*-/-*^) (**Fig. 2E**).

Considering the major changes in IMM ultrastructure in *Opa1*^*-/-*^ cells, we next looked at the nucleoid-cristae proximity, by immunofluorescence labeling of dsDNA and MICOS component Mic60. In WT MEFs, the nucleoids are abundant and interspaced in between the cristae as expected, displaying close mutual proximity (**Figure 2F**). However, in *Opa1*^*-/-*^ cells, there is an apparent disorganization of the nucleoid-cristae proximity. These data were further evaluated by analyzing the colocalization of mtDNA to Mic60 and vice versa. We found that mtDNA colocalization to Mic60 is decreased in *Opa1*^*-/-*^ cells and is mildly rescued after OPA1 stable expression, hence, nucleoid-cristae proximity is perturbed in mitochondria that contain nucleoids (**Figure 2F-G**). In contrast, Mic60 colocalization to mtDNA is rescued in *Opa1*^*-/-*^ cells after OPA1 expression, which relates to the mtDNA content and cristae periodicity recovery (**Figure 2F, H**).

Thus, in *Opa1*^*-/-*^ cells, mitochondria predominantly display sparse cristae with decreased nucleoid-cristae proximity, that are partly rescued by OPA1 stable expression.

### Acute expression of WT and pathogenic *OPA1* variants perturb mitochondrial nucleoid abundance and distribution

We next inspected the effect of OPA1 ADOA-causing mutants on nucleoid distribution and abundance to further understand what causes the mtDNA defects found in ADOA patients (18,19,46,47). We used our previously characterized model of overexpression of OPA1 in WT MEF cells and expression of two OPA1 ADOA-causing mutants (48,51): *OPA1* c.870+5G>A (p. Lys262_Arg290Del), located in the GTPase domain; and c.2713C>T (p.Arg905X), located in the GTPase-effector domain (GED) (**Fig. 3A**).

**Figure 3.**
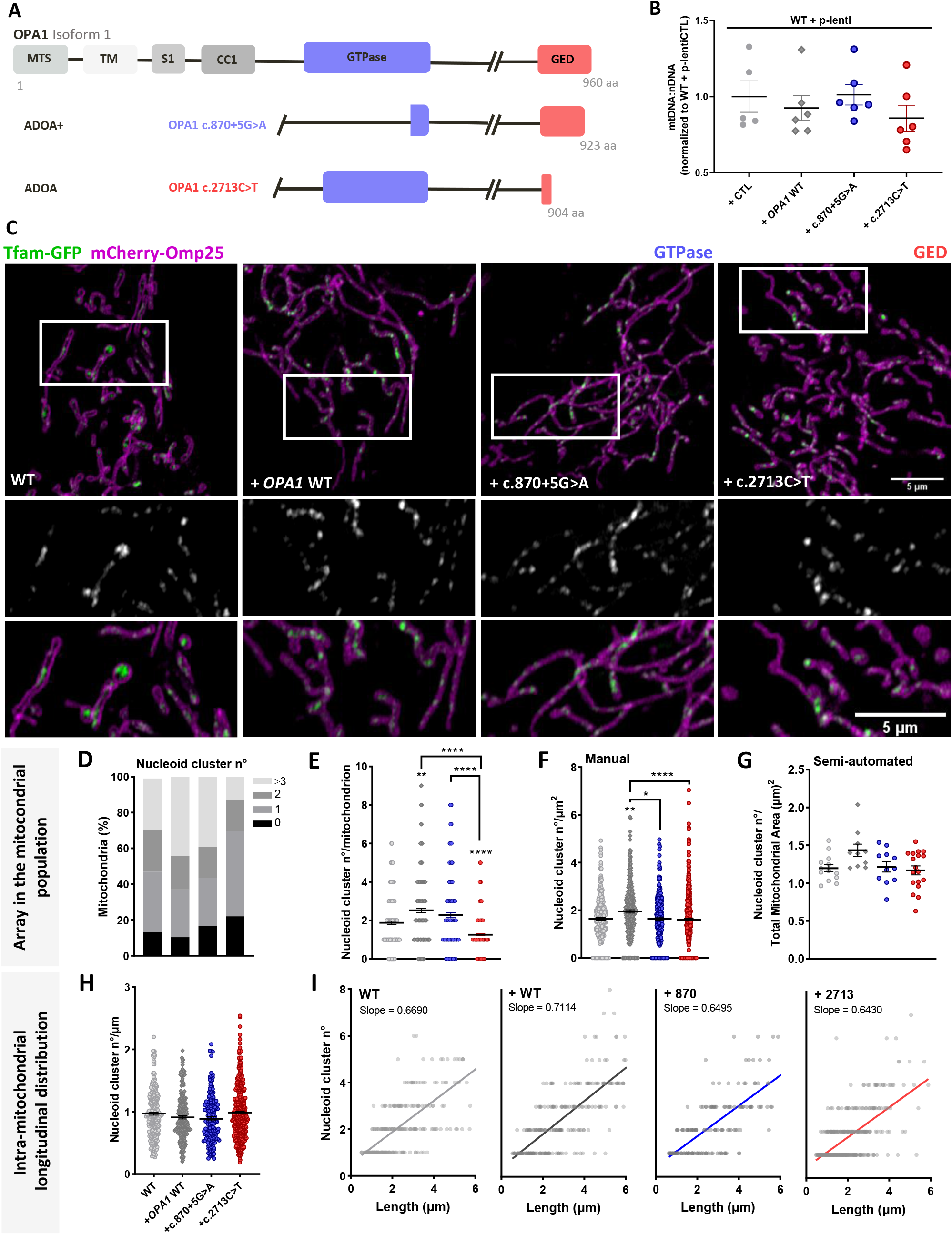
Acute expression of WT and pathogenic *OPA1* variants perturb mitochondrial nucleoid cluster abundance and distribution. **(A)** Schematic representation of OPA1 isoform 1 domains and ADOA-causing mutants: OPA1 c.870+5G>A (GTPase domain) and OPA1 c.2713C>T (GED). MTS: mitochondrial targeting sequence. TM: transmembrane domain. S1: Processing site 1. CC1: coiled-coil domain 1. **(B)** Mitochondrial DNA abundance of WT MEFs stably expressing a lentiviral plasmid carrying control cDNA (CTL), *OPA1* WT, or pathogenic *OPA1* variants; quantified by qPCR of mt-Nd4 and Gadph ratio. Data are mean ± SEM from ≥ 5 independent experiments. Scale bar = 1μm. **(C)** Representative images of WT MEFs, WT MEFs overexpressing *OPA1* WT, or pathogenic *OPA1* variants, co-transfected with mCherry-Omp25 (OMM) and Tfam-GFP (nucleoids) cDNA. Bottom panel, insets of Tfam-GFP (top), and merge (bottom). **(D)** Percentage of mitochondria with a different number of nucleoid clusters. **(E)** Mean nucleoid cluster number per mitochondrion. **(F)** Mean nucleoid cluster number per mitochondrion area manually quantified. **(G)** Mean nucleoid cluster number per total mitochondrial area, quantified semi-automatically by the MiNuD algorithm. **(H)** Mean nucleoid cluster number per mitochondrion length. **(I)** A simple regression of the relation between mitochondrial length and the number of nucleoids in single organelles. Data are mean ± SEM of ≥ 228 objects from ≥ 18 cells of ≥ 4 independent experiments. For G, data are mean ± SEM of ≥ 11 cells of ≥ 4 independent experiments (^****^p<0.0001; ^**^ p<0.005; ^*^ p<0.05).

We first analyzed mtDNA maintenance. Interestingly, acute, and stable overexpression of OPA1 showed no differences in mtDNA copy number. (**Fig. S5A; Fig. 3B)**. Likewise, mtDNA abundance was unchanged after stable overexpression of OPA1 mutants (**Fig. 3B**) or TFAM overexpression (**Fig. S5B)**. Moreover, the stable expression of OPA1 mutants did not cause mtDNA deletions (**Fig. S5C-D)**.

The nucleoid array showed that acute OPA1 overexpression caused a slight reduction in empty mitochondria (10% vs. 13% WT MEF) and an increase in the number of nucleoid clusters per mitochondrion and per mitochondrion area (**Fig 3C-G**). Conversely, cells with OPA1 c.870+5G>A expression showed an increase in “empty” mitochondria (17% vs. 10% WT OPA1) and a lower abundance of nucleoids clusters per mitochondrion area (**Fig. 3C-G**). Furthermore, cells expressing the mutant OPA1 c.2713C>T showed an increase in “empty” mitochondria (22% vs. 10% WT OPA1) and a reduction in the number of nucleoid clusters per mitochondrion and area (**Fig. 3C-G**). These changes were independent of the mitochondrial morphology since WT OPA1 and c.870+5G>A mutant overexpression showed elongated mitochondria, and cells expressing OPA1 c.2713C>T exhibited fragmented mitochondria (**Fig. S6A-D**), as we previously described in this model (48). Intramitochondrial longitudinal distribution of nucleoids was quantified in mitochondria up to 6 μm, the longest organelles found in OPA1 c.2713C>T (**Fig. S6C**). Interestingly, we found no changes in this parameter upon the expression of WT OPA1 or the studied mutants (**Fig. 3H-I)**.

Thus, in our model neither OPA1 overexpression nor the expression of ADOA-causing mutants induced changes in mtDNA abundance and integrity. Yet, OPA1 overexpression caused mitochondrial elongation and an increased nucleoid cluster abundance in the mitochondrial population. Surprisingly, cells expressing OPA1 ADOA-causing mutants showed alterations in mitochondrial morphology and nucleoid array in the mitochondrial population without altering the intramitochondrial longitudinal distribution of the nucleoids. Hence, this heterozygous-like system constitutes a pathologically relevant model to study nucleoid distribution upon alterations of mitochondrial dynamics without mtDNA depletion.

### Cristae ultrastructure defects in stable cell lines expressing *OPA1* ADOA-causing mutants

Previous data indicate that ADOA patients show IMM ultrastructure defects (40,48,52). Thus, we further evaluated the IMM ultrastructure. Our data show that stable OPA1 overexpression leads to an increase in short cristae (17% vs. 8% CTL) and aberrant cristae (23% vs. 16% CTL) (**Fig. 4A-B**). OPA1 c.870+5G>A mutant ‘s expression caused the same tendency, with an increase in short cristae (21% vs. 17% WT OPA1) and aberrant cristae (30% vs. 23% WT OPA1). Interestingly, cells expressing the mutant c.2713C>T showed similar abundance of aberrant cristae (21% vs. 23% WT OPA1) but a decrease in short cristae (8% vs. 17% WT OPA1) (**Fig. 4A-B**).

**Figure 4.**
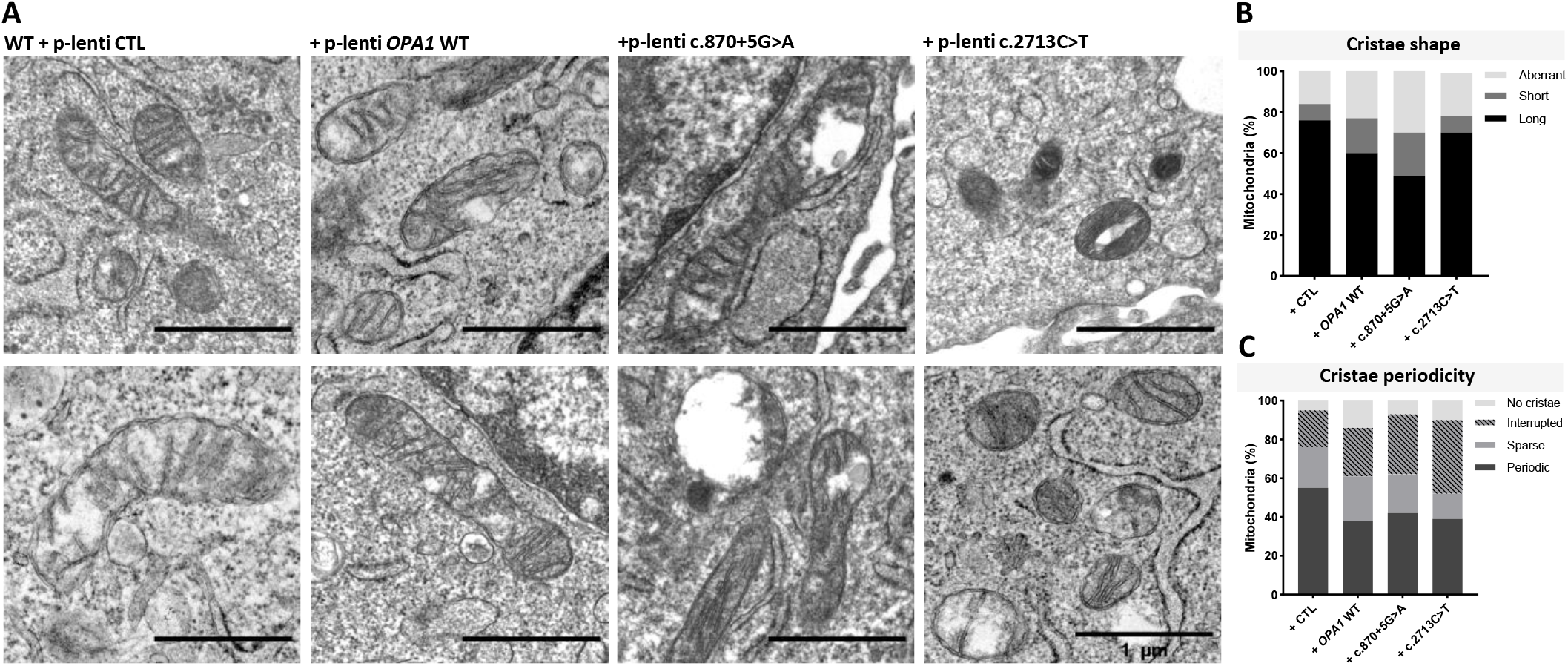
WT and pathogenic *OPA1* variant ‘s stable expression alter mitochondrial cristae ultrastructure. **(A)** Representative TEM images of WT MEFs stably expressing a lentiviral plasmid containing control cDNA (CTL), *OPA1* WT, or pathogenic *OPA1* variants. **(B)** Quantification of mitochondrial cristae shape descriptors. **(C)** Quantification of mitochondrial cristae periodicity. Data are from ≥ 112 objects from ≥ 3 independent experiments.

We then quantified cristae periodicity and found that cells with stable overexpression of OPA1 showed a decrease in periodic cristae (**Fig. 4A-C**). The expression of both mutants caused an increase in the interrupted cristae phenotype (31% c.870+5G>A; 38% c.2713C>T vs. 25% WT OPA1) (**Fig. 4A-C**). Therefore, OPA1 is relevant for cristae structuration and proper nucleoid array in the mitochondrial population. Nevertheless, we did not observe clear domain-specific differences of OPA1 mutants on nucleoid distribution and IMM structure.

## DISCUSSION

Here, we demonstrated that the absence of OPA1 or the expression of OPA1 ADOA-causing mutants, lead to an altered nucleoid array, suggesting that the execution of mitochondrial fusion is needed to keep nucleoid distribution amongst the mitochondrial population. Also, single-organelle analysis showed that the absence of OPA1 alters the cristae periodicity and nucleoids-to-cristae proximity, leading to perturbed longitudinal nucleoid distribution within each mitochondrion. Yet, the expression of OPA1 disease-causing mutants showed some defects in cristae periodicity, but no changes in longitudinal nucleoid distribution (**Fig. 5**). Thus, these data allow us to dissect the role of OPA1, with fusion supporting nucleoids array amongst organelles, while cristae shaping determining intra-mitochondrial mtDNA distribution.

**Figure 5.**
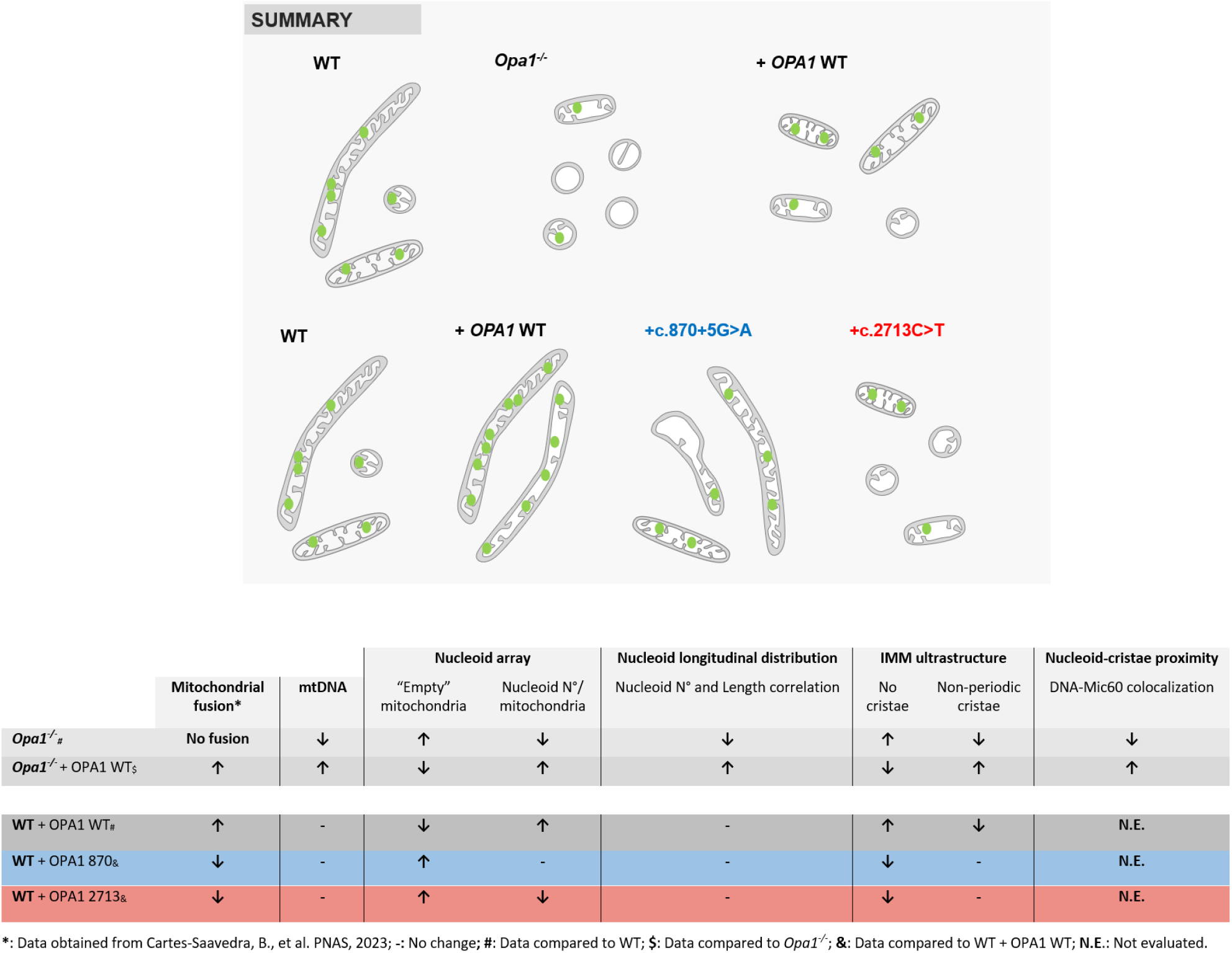
Summary of main findings. The cartoon and table show mitochondrial morphology and cristae ultrastructure changes, as well as nucleoid (green circles) abundance and nucleoid distribution alterations in the studied models. Mitochondrial fusion data were obtained from Cartes-Saavedra, et al. (48).

Different groups have studied the role of OPA1 in maintaining mtDNA and nucleoid abundance in mammalian cells (8,33,39,41,42,53). Yet, we emphasize the importance of describing nucleoid distribution considering the mitochondria available to contain them. Hence, we proposed a new paradigm based on our nucleoid distribution model (**Fig. 1B**), that incorporates mitochondrial characteristics and single-organelle analysis. Nucleoid distribution has gained attention as a functional parameter related to mtDNA distribution and inheritance (23,54,55), and cell function. Using this new paradigm, we showed that OPA1 plays a vital role in the distribution of the nucleoids in the mitochondrial population and at a single organelle level.

We found that in the absence of Opa1 and further rescue with OPA1 isoform 1, there is a correlation between mitochondrial morphology, cristae ultrastructure, mtDNA levels, and nucleoid number, consistent with previous reports (41). Thus, the rescue of OPA1 isoform 1 restores mitochondrial fusion, morphology and cristae ultrastructure supporting the semi-regular distribution of the nucleoids and mtDNA levels, independently from the direct interaction proposed between OPA1exon 4b isoforms and mtDNA (39,42). Previous studies showed that rescue of the mtDNA levels upon the expression of different OPA1 isoforms in *Opa1*^*-/-*^ cells are related to rescue of fusion and IMM ultrastructure and not to morphology changes (41), highlighting these two components as possible regulators of mtDNA abundance.

OPA1 overexpression increased mitochondrial fusion rate (48), but the relative abundance of mtDNA molecules was unaltered, however, the nucleoids are more spread within the mitochondrial population than in WT cells. This supports the idea that fusion-induced mitochondrial elongation or material exchange play a role in the distribution of nucleoids amongst mitochondria, as has been showed in the Mfn 1/2 KO model (56).

We did not find mtDNA deletions in our stable cell lines with mutant OPA1, although multiple mtDNA deletions have been found in skeletal muscle biopsies from ADOA patients (46). The accumulation of mtDNA damage is characteristic for post-mitotic cells and it emphasizes the relevance of studying nucleoid distribution in post-mitotic tissues and on subcellular level, as thelocal accumulation of damaged mtDNA causes bioenergetic dysfunction in these patients (2,18).

The nucleoid array in the mitochondrial network describes how they are spread, which is relevant for the local bioenergetic function of individual mitochondria and the cells. It is proposed that mtDNA has a “sphere of influence” over the nearby cristae, locally controlling OXPHOS function (57,58). Since mitochondria without nucleoids (“empty”) are accumulated in *Opa1*^*-/-*^ cells and WT cells expressing the ADOA-mutants, this phenotype might be related to the overall bioenergetic dysfunction found in these cells (51,59), and most importantly, in ADOA patients (18). In WT MEFs overexpressing OPA1 and the c.2713C>T ADOA-causing mutant, nucleoid array was altered on the organelle ‘s population, associated with cristae disorganization. However, expression of the c.870+5G>A mutant caused cristae alterations but a milder effect on the nucleoid array in the mitochondrial population, suggesting that nucleoid distribution amongst mitochondria is not exclusively directed by cristae ultrastructure.

Interestingly, intramitochondrial longitudinal distribution of nucleoids was a robust parameter only altered in *Opa1*^*-* /-^ cells, that loss cristae periodicity and nucleoids-to-cristae proximity. Both parameters were partially rescued after OPA1 isoform 1 expression, suggesting a key role of this isoform in the arrangement of nucleoids within each mitochondrion, and the periodicity of IMM folding. This finding aligns with the super-resolution data that show the nucleoid interspersed between the cristae in live cells (60) and the nucleoid disorganization upon cristae disorganization due to Mic60 silencing (61).

In conclusion, our new paradigm allowed us to characterize nucleoid distribution in relation to the characteristics of the mitochondria that contain them. We showed that isoform 1 of OPA1 plays a vital role in the distribution of the nucleoids in the mitochondrial population and single organelle level by means of IMM fusion and cristae organization. Furthermore, OPA1 disease-causing mutants lead to altered nucleoid distribution irrespective of the protein domain where the mutation is located, associated with fusion inhibition and cristae morphology defects, providing new insights into the pathophysiology of ADOA.

## MATERIALS AND METHODS

Extended Materials and Methods can be found in Supplementary Information.

### Live-cell nucleoid cluster imaging

MEF cells were plated on 25 mm glass coverslips and grown in culture up to 80% confluence. Nucleoids stained using 1μL/mL PicoGreen (15 min, 37°C; Thermo Fisher Scientific) or transfection with Tfam-GFP. Mitochondria were stained with MitoTracker Deep Red 75nM (15 min, 37°C; Thermo Fisher Scientific) or transfection of mCherry-Omp25. Live-cell imaging was performed in a 0.25% BSA extracellular medium at 37°C as previously described (51). Image acquisition was performed with 5 Z-stacks (0,19 μm interval) in a confocal microscope Zeiss LSM 880 with an Airyscan detector (63X PlanApo 1.4 NA) on the Fast SR mode, at the Advanced Microscopy Facility UMA UC. After acquisition, images were deconvoluted in 3D using the ZEN Black software.

### Nucleoid cluster distribution quantification

We manually quantified nucleoid distribution and abundance inside single organelles (**Fig. S1A**). The manual analysis was performed in FIJI ImageJ. We detected a high Tfam-GFP intramitochondrial background noise *Opa1*^*-/-*^ cells, previously described as “diffused TFAM” (39). Thus, we adjusted the B/C settings to distinguish the brightest puncta (**Fig. S3A-C**). To strengthen our distribution data in the mitochondrial population, we developed a semi-automated method as a Python package (https://github.com/RudgeLab/MiNuD) (**Fig. S1B**). The input for MiNuD is the .tif files of light-microscopy images of mitochondria, nucleoids, and their respective masks obtained from FIJI threshold or Trainable Weka.

## ACKNOWLEDGMENTS

We thank Drs. Karin Busch, Marco Tigano and György Hajnóczky for thoughtful discussion; Alejandro Munizaga and Liseth Garibaldi for TEM sample preparation, Nicole Salgado, and Fernanda Gárate for imaging technical assistance at the Advanced Microscopy Facility at Pontificia Universidad Católica de Chile. The Chilean Government supported this work through Agencia Nacional de Investigación y Desarrollo (ANID) Ph.D. Fellowships 21211363 to JM, 21201041 to IM-R, 21191304 to DL, and 21181402 to BC-S. GV was supported by scholarships from the Institute for Biological and Medical Engineering, Pontificia Universidad Católica de Chile, and the School of Computing, Newcastle University. ANID PIA/ACT192015 to TJR. FONDECYT grants 1191770 to VE and TJR, 1211598 to TJR, and 1231557 to VE. R.H. is a Wellcome Trust Investigator (109915/Z/15/Z), who received support from the Medical Research Council (UK) (MR/V009346/1), the Newton Fund (UK/Turkey, MR/N027302/1), the Addenbrookes Charitable Trust (G100142), the Evelyn Trust, the Stoneygate Trust, the Lily Foundation, Ataxia UK, Action for AT and an MRC strategic award to establish an International Centre for Genomic Medicine in Neuromuscular Diseases (ICGNMD) MR/S005021/1. This research was supported by the NIHR Cambridge Biomedical Research Centre (BRC-1215-20014). The views expressed are those of the authors and not necessarily those of the NIHR or the Department of Health and Social Care.

## REFERENCES

1. zzEisner V, Picard M, Hajnóczky G. Mitochondrial dynamics in adaptive and maladaptive cellular stress responses. Nature Cell Biology. 2018 Jul;20(7):755–65.

2. Vincent AE, Rosa HS, Pabis K, Lawless C, Chen C, Grünewald A, et al. Subcellular origin of mitochondrial DNA deletions in human skeletal muscle. Annals of Neurology. 2018;84(2):289–301.

3. Yu-Wai-Man P, Lai-Cheong J, Borthwick GM, He L, Taylor GA, Greaves LC, et al. Somatic Mitochondrial DNA Deletions Accumulate to High Levels in Aging Human Extraocular Muscles. Investigative Ophthalmology & Visual Science. 2010 Jul 1;51(7):3347–53.

4. Brown TA, Tkachuk AN, Shtengel G, Kopek BG, Bogenhagen DF, Hess HF, et al. Superresolution Fluorescence Imaging of Mitochondrial Nucleoids Reveals Their Spatial Range, Limits, and Membrane Interaction. Molecular and Cellular Biology. 2011 Dec 15;31(24):4994–5010.

5. Kukat C, Wurm CA, Spåhr H, Falkenberg M, Larsson NG, Jakobs S. Super-resolution microscopy reveals that mammalian mitochondrial nucleoids have a uniform size and frequently contain a single copy of mtDNA. Proc Natl Acad Sci USA. 2011 Aug 16;108(33):13534–9.

6. Kukat C, Davies KM, Wurm CA, Spåhr H, Bonekamp NA, Kühl I, et al. Cross-strand binding of TFAM to a single mtDNA molecule forms the mitochondrial nucleoid. PNAS. 2015 Sep 8;112(36):11288–93.

7. Ngo H, Jt K, Dc C. The mitochondrial transcription and packaging factor Tfam imposes a U-turn on mitochondrial DNA. Nat Struct Mol Biol. 2011 Oct 30;18(11):1290–6.

8. Chen H, Vermulst M, Wang YE, Chomyn A, Prolla TA, McCaffery JM, et al. Mitochondrial Fusion Is Required for mtDNA Stability in Skeletal Muscle and Tolerance of mtDNA Mutations. Cell. 2010 Apr 16;141(2):280–9.

9. Ishihara T, Ban-Ishihara R, Maeda M, Matsunaga Y, Ichimura A, Kyogoku S, et al. Dynamics of Mitochondrial DNA Nucleoids Regulated by Mitochondrial Fission Is Essential for Maintenance of Homogeneously Active Mitochondria during Neonatal Heart Development. Molecular and Cellular Biology. 2015 Jan 1;35(1):211–23.

10. Ban-Ishihara R, Ishihara T, Sasaki N, Mihara K, Ishihara N. Dynamics of nucleoid structure regulated by mitochondrial fission contributes to cristae reformation and release of cytochrome c. Proc Natl Acad Sci USA. 2013 Jul 16;110(29):11863–8.

11. Ota A, Ishihara T, Ishihara N. Mitochondrial nucleoid morphology and respiratory function are altered in Drp1-deficient HeLa cells. J Biochem. 2020 Mar 1;167(3):287–94.

12. Tezze C, Romanello V, Desbats MA, Fadini GP, Albiero M, Favaro G, et al. Age-Associated Loss of OPA1 in Muscle Impacts Muscle Mass, Metabolic Homeostasis, Systemic Inflammation, and Epithelial Senescence. Cell Metab. 2017 Jun 6;25(6):1374–1389.e6.

13. Renaldo F, Amati-Bonneau P, Slama A, Romana C, Forin V, Doummar D, et al. MFN2, a new gene responsible for mitochondrial DNA depletion. Brain. 2012 Aug 1;135(8):e223–e223.

14. Rouzier C, Bannwarth S, Chaussenot A, Chevrollier A, Verschueren A, Bonello-Palot N, et al. The MFN2 gene is responsible for mitochondrial DNA instability and optic atrophy ‘plus ‘ phenotype. Brain. 2012 Jan 1;135(1):23–34.

15. Bannwarth S, Ait-El-Mkadem S, Chaussenot A, Genin EC, Lacas-Gervais S, Fragaki K, et al. A mitochondrial origin for frontotemporal dementia and amyotrophic lateral sclerosis through CHCHD10 involvement. Brain. 2014 Aug 1;137(8):2329–45.

16. Genin EC, Plutino M, Bannwarth S, Villa E, Cisneros-Barroso E, Roy M, et al. CHCHD10 mutations promote loss of mitochondrial cristae junctions with impaired mitochondrial genome maintenance and inhibition of apoptosis. EMBO Molecular Medicine. 2016 Jan 1;8(1):58–72.

17. Vielhaber S, Debska-Vielhaber G, Peeva V, Schoeler S, Kudin AP, Minin I, et al. Mitofusin 2 mutations affect mitochondrial function by mitochondrial DNA depletion. Acta Neuropathol. 2013 Feb;125(2):245–56.

18. Yu-Wai-Man P, Sitarz KS, Samuels DC, Griffiths PG, Reeve AK, Bindoff LA, et al. OPA1 mutations cause cytochrome c oxidase deficiency due to loss of wild-type mtDNA molecules. Hum Mol Genet. 2010 Aug 1;19(15):3043–52.

19. Sitarz KS, Almind GJ, Horvath R, Czermin B, Grønskov K, Pyle A, et al. OPA1 mutations induce mtDNA proliferation in leukocytes of patients with dominant optic atrophy. Neurology. 2012 Oct 2;79(14):1515–7.

20. Ono T, Isobe K, Nakada K, Hayashi JI. Human cells are protected from mitochondrial dysfunction by complementation of DNA products in fused mitochondria. Nature Genetics. 2001 Jul;28(3):272–5.

21. Iborra FJ, Kimura H, Cook PR. The functional organization of mitochondrial genomes in human cells. BMC Biology. 2004 May 24;2(1):9.

22. Legros F, Malka F, Frachon P, Lombès A, Rojo M. Organization and dynamics of human mitochondrial DNA. Journal of Cell Science. 2004 Jun 1;117(13):2653–62.

23. Jajoo R, Jung Y, Huh D, Viana MP, Rafelski SM, Springer M, et al. Accurate concentration control of mitochondria and nucleoids. Science. 2016 Jan 8;351(6269):169–72.

24. Lewis SC, Uchiyama LF, Nunnari J. ER-mitochondria contacts couple mtDNA synthesis with mitochondrial division in human cells. Science. 2016 Jul 15;353(6296):aaf5549.

25. Jakobs S, Stephan T, Ilgen P, Brüser C. Light Microscopy of Mitochondria at the Nanoscale. Annu Rev Biophys. 2020 May 6;49(1):289–308.

26. Dlasková A, Engstová H, Špaček T, Kahancová A, Pavluch V, Smolková K, et al. 3D super-resolution microscopy reflects mitochondrial cristae alternations and mtDNA nucleoid size and distribution. Biochimica et Biophysica Acta (BBA) - Bioenergetics. 2018 Sep 1;1859(9):829–44.

27. Kopek BG, Shtengel G, Xu CS, Clayton DA, Hess HF. Correlative 3D superresolution fluorescence and electron microscopy reveal the relationship of mitochondrial nucleoids to membranes. PNAS. 2012 Apr 17;109(16):6136–41.

28. Blum TB, Hahn A, Meier T, Davies KM, Kühlbrandt W. Dimers of mitochondrial ATP synthase induce membrane curvature and self-assemble into rows. PNAS. 2019 Mar 5;116(10):4250–5.

29. Stephan T, Brüser C, Deckers M, Steyer AM, Balzarotti F, Barbot M, et al. MICOS assembly controls mitochondrial inner membrane remodeling and crista junction redistribution to mediate cristae formation. The EMBO Journal. 2020 Jul 15;39(14):e104105.

30. Kondadi AK, Anand R, Hänsch S, Urbach J, Zobel T, Wolf DM, et al. Cristae undergo continuous cycles of membrane remodelling in a MICOS-dependent manner. EMBO reports. 2020 Mar 4;21(3):e49776.

31. Quintana-Cabrera R, Quirin C, Glytsou C, Corrado M, Urbani A, Pellattiero A, et al. The cristae modulator Optic atrophy 1 requires mitochondrial ATP synthase oligomers to safeguard mitochondrial function. Nature Communications. 2018 Aug 24;9(1):3399.

32. Frezza C, Cipolat S, Brito OM de, Micaroni M, Beznoussenko GV, Rudka T, et al. OPA1 Controls Apoptotic Cristae Remodeling Independently from Mitochondrial Fusion. Cell. 2006 Jul 14;126(1):177–89.

33. Fry MY, Navarro PP, Hakim P, Ananda VY, Qin X, Landoni JC, et al. In situ architecture of Opa1-dependent mitochondrial cristae remodeling. The EMBO Journal. 2024 Jan 15;1–23.

34. Song Z, Ghochani M, McCaffery JM, Frey TG, Chan DC. Mitofusins and OPA1 Mediate Sequential Steps in Mitochondrial Membrane Fusion. Mol Biol Cell. 2009 Aug 1;20(15):3525–32.

35. Liu X, Weaver D, Shirihai O, Hajnóczky G. Mitochondrial ‘kiss-and-run ‘: interplay between mitochondrial motility and fusion–fission dynamics. The EMBO Journal. 2009 Oct 21;28(20):3074–89.

36. Anand R, Wai T, Baker MJ, Kladt N, Schauss AC, Rugarli E, et al. The i-AAA protease YME1L and OMA1 cleave OPA1 to balance mitochondrial fusion and fission. J Cell Biol. 2014 Mar 17;204(6):919–29.

37. Herlan M, Vogel F, Bornhovd C, Neupert W, Reichert AS. Processing of Mgm1 by the rhomboid-type protease Pcp1 is required for maintenance of mitochondrial morphology and of mitochondrial DNA. J Biol Chem. 2003 Jul 25;278(30):27781–8.

38. Sesaki H, Southard SM, Yaffe MP, Jensen RE. Mgm1p, a Dynamin-related GTPase, Is Essential for Fusion of the Mitochondrial Outer Membrane. MBoC. 2003 Jun;14(6):2342–56.

39. Yang L, Tang H, Lin X, Wu Y, Zeng S, Pan Y, et al. OPA1-Exon4b Binds to mtDNA D-Loop for Transcriptional and Metabolic Modulation, Independent of Mitochondrial Fusion. Front Cell Dev Biol. 2020;8.

40. Patten DA, Wong J, Khacho M, Soubannier V, Mailloux RJ, Pilon-Larose K, et al. OPA1-dependent cristae modulation is essential for cellular adaptation to metabolic demand. The EMBO Journal. 2014 Nov 18;33(22):2676–91.

41. Del Dotto V, Mishra P, Vidoni S, Fogazza M, Maresca A, Caporali L, et al. OPA1 Isoforms in the Hierarchical Organization of Mitochondrial Functions. Cell Reports. 2017 Jun 20;19(12):2557–71.

42. Elachouri G, Vidoni S, Zanna C, Pattyn A, Boukhaddaoui H, Gaget K, et al. OPA1 links human mitochondrial genome maintenance to mtDNA replication and distribution. Genome Res. 2011 Jan 1;21(1):12–20.

43. Lenaers G, Hamel C, Delettre C, Amati-Bonneau P, Procaccio V, Bonneau D, et al. Dominant optic atrophy. Orphanet Journal of Rare Diseases. 2012 Jul 9;7(1):46.

44. Yu-Wai-Man P, Trenell MI, Hollingsworth KG, Griffiths PG, Chinnery PF. OPA1 mutations impair mitochondrial function in both pure and complicated dominant optic atrophy. Brain. 2011 Apr 1;134(4):e164–e164.

45. Yu-Wai-Man P, Griffiths PG, Gorman GS, Lourenco CM, Wright AF, Auer-Grumbach M, et al. Multi-system neurological disease is common in patients with OPA1 mutations. Brain. 2010 Mar 1;133(3):771–86.

46. Amati-Bonneau P, Valentino ML, Reynier P, Gallardo ME, Bornstein B, Boissière A, et al. OPA1 mutations induce mitochondrial DNA instability and optic atrophy ‘plus ‘ phenotypes. Brain. 2008 Feb 1;131(2):338–51.

47. Hudson G, Amati-Bonneau P, Blakely EL, Stewart JD, He L, Schaefer AM, et al. Mutation of OPA1 causes dominant optic atrophy with external ophthalmoplegia, ataxia, deafness and multiple mitochondrial DNA deletions: a novel disorder of mtDNA maintenance. Brain. 2008 Feb 1;131(2):329–37.

48. Cartes-Saavedra B, Lagos D, Macuada J, Arancibia D, Burté F, Sjöberg-Herrera MK, et al. OPA1 disease-causing mutants have domain-specific effects on mitochondrial ultrastructure and fusion. Proceedings of the National Academy of Sciences. 2023 Mar 21;120(12):e2207471120.

49. Pastukh V, Shokolenko I, Wang B, Wilson G, Alexeyev M. Human mitochondrial transcription factor A possesses multiple subcellular targeting signals. The FEBS Journal. 2007 Dec 1;274(24):6488–99.

50. Huff J. The Fast mode for ZEISS LSM 880 with Airyscan: high-speed confocal imaging with super-resolution and improved signal-to-noise ratio. Nature Methods. 2016 Nov;13(11):i–ii.

51. Cartes-Saavedra B, Macuada J, Lagos D, Arancibia D, Andrés ME, Yu-Wai-Man P, et al. OPA1 Modulates Mitochondrial Ca2+ Uptake Through ER-Mitochondria Coupling. Frontiers in Cell and Developmental Biology. 2022;9.

52. Agier V, Oliviero P, Lainé J, L’Hermitte-Stead C, Girard S, Fillaut S, et al. Defective mitochondrial fusion, altered respiratory function, and distorted cristae structure in skin fibroblasts with heterozygous OPA1 mutations. Biochimica et Biophysica Acta (BBA) - Molecular Basis of Disease. 2012 Oct 1;1822(10):1570–80.

53. Rodríguez-Nuevo A, Díaz-Ramos A, Noguera E, Díaz-Sáez F, Duran X, Muñoz JP, et al. Mitochondrial DNA and TLR9 drive muscle inflammation upon Opa1 deficiency. The EMBO Journal. 2018 May 15;37(10):e96553.

54. Tauber J, Dlasková A, Šantorová J, Smolková K, Alán L, Špaček T, et al. Distribution of mitochondrial nucleoids upon mitochondrial network fragmentation and network reintegration in HEPG2 cells. The International Journal of Biochemistry & Cell Biology. 2013 Mar 1;45(3):593–603.

55. Ilamathi HS, Ouellet M, Sabouny R, Desrochers-Goyette J, Lines MA, Pfeffer G, et al. A new automated tool to quantify nucleoid distribution within mitochondrial networks. Sci Rep. 2021 Nov 23;11(1):22755.

56. Silva Ramos E, Motori E, Brüser C, Kühl I, Yeroslaviz A, Ruzzenente B, et al. Mitochondrial fusion is required for regulation of mitochondrial DNA replication. Barsh GS, editor. PLoS Genet. 2019 Jun 6;15(6):e1008085.

57. Busch KB, Kowald A, Spelbrink JN. Quality matters: how does mitochondrial network dynamics and quality control impact on mtDNA integrity? Philosophical Transactions of the Royal Society B: Biological Sciences. 2014 Jul 5;369(1646):20130442.

58. Jakubke C, Roussou R, Maiser A, Schug C, Thoma F, Bunk D, et al. Cristae-dependent quality control of the mitochondrial genome. Science Advances. 2021;7(36):eabi8886.

59. Olichon A, Landes T, Arnauné-Pelloquin L, Emorine LJ, Mils V, Guichet A, et al. Effects of OPA1 mutations on mitochondrial morphology and apoptosis: Relevance to ADOA pathogenesis. Journal of Cellular Physiology. 2007;211(2):423–30.

60. Liu T, Stephan T, Chen P, Keller-Findeisen J, Chen J, Riedel D, et al. Multi-color live-cell STED nanoscopy of mitochondria with a gentle inner membrane stain. Proc Natl Acad Sci U S A. 2022 Dec 27;119(52):e2215799119.

61. Li H, Ruan Y, Zhang K, Jian F, Hu C, Miao L, et al. Mic60/Mitofilin determines MICOS assembly essential for mitochondrial dynamics and mtDNA nucleoid organization. Cell Death & Differentiation. 2016 Mar;23(3):380–92.

62. Young L, Sung J, Stacey G, Masters JR. Detection of Mycoplasma in cell cultures. Nat Protoc. 2010 May;5(5):929–34.

63. Csordás G, Várnai P, Golenár T, Roy S, Purkins G, Schneider TG, et al. Imaging interorganelle contacts and local calcium dynamics at the ER-mitochondrial interface. Mol Cell. 2010 Jul 9;39(1):121–32.

