## Supplementary material for "*OPA1* and disease-causing mutants perturb mitochondrial nucleoid distribution"

### SUPPLEMENTARY INFORMATION

### EXTENDED MATERIALS AND METHODS

#### Cell culture

Experiments were performed in WT and *Opa1*$^{-/-}$ Murine embryonic fibroblasts (MEFs), kindly donated by Dr. György Hajnóczky. All the cells were cultured in high glucose Dulbecco-Eagle modified medium containing sodium pyruvate (DMEM, Gibco Cat #1280017) and supplemented with 10% of fetal bovine serum (FBS, Sigma Cat #F0926), 2mM GlutaMAX (Gibco, Cat #35050061), 100 U/ml Penicillin and 100 µg/mL Streptomycin (Gibco, Cat #15140-122), and 3,7 g/L Sodium Bicarbonate ($NaHCO_3$, Sigma-Aldrich). Cells were maintained at 37ºC in humidified air with 5% $CO_2$. All the cells were tested for mycoplasma contamination regularly using the following protocol (62). The cells were cultured following institutional bioethics protocols.

#### Cell Transfection

Cells were plated on 35 mm dishes with glass 25 mm coverslips and then transfected with specific constructs. MEF cells were transfected with a pCCEY plasmid encoding for human OPA1 isoform 1, kindly donated by Dr. Guy Lenaers. OPA1 GTPase or GED mutants were built in the same backbone as previously described (51). For nucleoid distribution analysis, we used a mCherry-Omp25 plasmid to label the OMM, and nucleoids were labeled with Tfam-GFP, kindly donated by Dr. György Hajnóczky and Dr. Jodi Nunnari, respectively. The transfection was performed with Opti-MEM (Gibco, Cat #51985-034) or Transfectagro (Corning, Cat #40-300-CVR) and Lipofectamine 3000 (Invitrogen, Cat#L300015), according to the manufacturer´s protocol. For optimal expression, the cells were grown for 48 h after transfection. Each experiment used one µg of DNA per plasmid per 35 mm dish.

#### Stable cell line generation

WT *or Opa1*$^{-/-}$ MEF cells were transfected as described above with a lentiviral plasmid containing the human OPA1 isoform 1 WT, c.870+5G>A or c.2713C>T mutation and a puromycin resistance cassette (pLentiOPA1, VectorBuilder). Transfected cells were selected with Puromycin 2 µg/mL to obtain a polyclonal stable cell line or further selected to obtain a monoclonal stable cell line, as previously described (48,51). The cells were grown in the same conditions described above in the presence of Puromycin 2 µg/mL.

#### Manual quantification of nucleoid cluster distribution in the mitochondrial population and intramitochondrial distribution

We established a manual method to quantify nucleoid distribution and abundance inside single organelles (**Fig. S1A**). We developed a complementary semi-automated method to strengthen our distribution data in the mitochondrial population (**Fig. S1B**). The manual analysis was performed in FIJI ImageJ as follows. We applied a

Z-projection in FIJI ImageJ and separated the two channels. We adjusted B/C settings in the mCherry-Omp25/MitoTracker Deep Red channel and selected recognizable signals as individual objects (i.e. not overlapped), without complex shapes (i.e. donuts), and that were entirely inside the field of view. The perimeter of each object was then manually drawn using the “freehand” tool. The length of each object was calculated by drawing a line inside the mitochondrion from tip-to-tip, using the “freehand” tool. Next, using “measure” we calculated each mitochondrion’s area, circularity, and length. These measurements were then associated with the number of Tfam-GFP/Picogreen foci >150 nm detected inside the perimeter of each mitochondrion to calculate the n° nucleoid clusters per mitochondrion, per mitochondrial area, and length.

On the *Opa1*$^{-/-}$ cells, we detected a high Tfam-GFP intramitochondrial background noise, previously described as “diffused TFAM” (39). To deal with this issue, we manually increased the B/C settings adjustment till we could distinguish only the brightest puncta. Applying the adjustment, we could visualize Tfam-GFP signals closely resembling PicoGreen and anti-DNA staining (**Fig. S4A-C**). Using these alternative nucleoid labeling methods, we could validate the decrease in nucleoid signals found in adjusted Tfam-GFP labeled samples previously reported for *Opa1*$^{-/-}$ cells (**Fig. S4A-C**) (39).

**Semi-automated analysis of mitochondrial and nucleoid cluster populations, MiNuD**

We developed a semi-automated method as a Python package (https://github.com/RudgeLab/MiNuD) to analyze light-microscopy data and quantify the nucleoid array in the mitochondria population in a large set of data, customizable to the needs of the user. The input for MiNuD is the .tif files of light-microscopy images of mitochondria, nucleoids, and their respective masks obtained from FIJI threshold or Trainable Weka. The MiNuD package is a customizable code that consists of a single class called *mito_segmentator* which has all the functionalities as “methods”. In the *mito_segmentator*, the user can select the threshold for mitochondria images, nucleoid images, and external segmentations. The default thresholds are Otsu for mitochondria and external segmentation; Triangle for nucleoids. Nevertheless, any threshold from Sci-kit image can be used. Additionally, MiNuD provides an option to select a minimal area for mitochondria and a minimal area for nucleoids. Smaller objects will be removed and not analyzed. It can also remove mitochondria that are not overlapped with any nucleoid and nucleoids that are not overlapped with any mitochondria, or both, to study mitochondria with nucleoids or empty mitochondria specifically. Additionally, it can remove objects touching the edges of the image and therefore are not entirely in the field of view.

The *mito_segmentator* can use the method *folder_analysis* to analyze many files automatically. It uses a directory as input and reads all the folders found inside. In the folders, MiNuD looks for patterns in the names of the files, which can be specified by the user, to get images of mitochondria and nucleoids for analysis. Then the areas for mitochondria and nucleoids are computed, converting pixels to μm$^2$. This information is shown in a *raw_data* file.

The *raw_data* includes the frame number (if the input is an image sequence), the applied thresholds, and the area and number of mitochondria and nucleoids found in that frame by the software. Furthermore, another output is the *analyzed data* file, which includes helpful information such as the number of nucleoids per mitochondrial area. The *raw_data* and *analyzed_data* can be obtained in XLSX, JSON, and Pandas. Finally, for visual inspection and quality check of the masks created by the semi-automated workflow, MiNuD has the method *get_masks*, which generates a superimposed image of the original mitochondrial image and its segmentation shown as red overlays, and the original nucleoid image and its segmentation shown as green overlays.

**Transmission Electron Microscopy**

For the ultrastructure studies, we used pellets of ≥ 8 × 105 cells, fixed with glutaraldehyde 2.5%. Staining was performed as previously described (63). Images of ultra-thin sections were acquired in a transmission electron microscope Philips Tecnai 12 at 80 kV or a TALOS F200C G2 system (Thermo Scientific) equipped with a Ceta 16M CMOS camera, at 200 kV, at the Advanced Microscopy Facility UMA UC, Pontificia Universidad Católica de Chile. The images were then analyzed, and each mitochondrion was classified regarding their cristae shape (long, short, and aberrant) and the distribution of their cristae, considered as the overall periodicity between cristae (periodic, non-periodic, no cristae). Each mitochondrion was included in only one category considering their main phenotype. If an aberrant crista was present in the mitochondrion, it was classified as aberrant, regardless of the abundance of long and short cristae. Non-periodic cristae were subclassified as interrupted and sparsed. The interrupted phenotype was characterized as a mitochondrion with a region of periodic cristae abruptly interrupted by an empty space. The sparse phenotype is characterized as a mitochondrion with few visible cristae, which are usually scattered and not periodically arranged. Short and long cristae were classified as previously described (48).

**Mitochondrial DNA quantification**

Total DNA from MEF cells was isolated with Wizard® Genomic DNA Purification kit (A1120, Promega), following the manufacturer's instructions. Quantitative real-time PCR was performed in triplicate in 96-well plates in a QuantStudio 3 thermocycler (Applied Biosystems) using KAPA SYBR FAST qPCR Master Mix (KK4601, Sigma Aldrich) or PowerUp™ SYBR™ Green Master (Applied Biosystems), and 10 ng of total DNA. The relative abundance of the mitochondrial DNA encoded gene mtNd4 relative to the nuclear DNA encoded gene Gapdh was determined according to the 2-ΔΔCt method and expressed as mtDNA:nDNA ratio. Primers sets for mtNd4: Forward Primer 5'AGCTCAATCTGCTTACGCCA3'; Reverse primer 5' TGTGAGGCCATGTGCGATTA 3'. Primers sets for Gadph: Forward primer 5'ATTTGCCGTGAGTGGAGTCATA 3', Reverse primer 5' CACTACAGACCCATGAGGAGTT 3'.

**Immunofluorescence analysis**

WT or *Opa1*$^{-/-}$ MEF cells were seeded in 15 mm glass coverslips and grown in culture up to 80% confluence. Cells were incubated for 45 min at 37°C with 150 nM MitoTracker Deep Red (Thermo Fisher Scientific) prior to fixation. Fixation was performed with 4% paraformaldehyde. (Thermo Fisher Scientific) for 15 min at room temperature and washed three times with cold PBS, permeabilized with 0.5% Triton X-100 for 10 min and blocked with 3% BSA (NC9418785, Rockland) for 1 hr at room temperature. The cells were incubated with the primary antibodies anti-DNA mouse monoclonal 1:50 (AC-30-10, Progen) and anti-Mic60 rabbit 1:250 or 1:1000 (10179-1-AP, Proteintech) overnight at 4 °C in a humid chamber. After washing three times with PBS, cells were incubated for 1hr at room temperature with 1:500 secondary antibody goat anti-rabbit IgG (H + L) highly cross-adsorbed labeled with Alexa Fluor 633 (Thermo Fisher Scientific) and 1:500 goat anti-mouse IgG (H + L) highly cross-adsorbed labeled with Alexa Fluor 568 (Thermo Fisher Scientific). Image acquisition was performed with 5 Z-stacks (0,17 µm interval) in a confocal microscope Zeiss LSM 880 with an Airyscan detector (63X PlanApo 1.4 NA), located at the Advanced Microscopy Facility UMA UC. For colocalization analysis, the images were cropped to avoid nuclei dsDNA signal. Colocalization coefficients were calculated in the software ZEN Blue as follows:

Colocalization coefficient channel 1 to channel 2= (#pixels$_{\text{channel 1}}$ colocalized/ #pixels$_{\text{channel 1}}$ total)

**Long-Range PCR**

MtDNA deletions were studied by Southern blot and/or long-range PCR using PrimeSTAR GXL polymerase (Takara Bio Europe) and 100 ng of DNA. The reaction was performed overnight for 30 cycles. Primer sets for mtDNA_7.7kb: Forward primer 5' ACCTGAATTGGGGGCCAACC 3'; Reverse primer 5' TGGCGAAGTGGGCTTTTGCT 3'. Primer sets for mtDNA_8.6kb: Forward primer 5' AGCAAAAGCCCACTTCGCCA 3'; Reverse primer 5' GGTTGGCCCCCAATTCAGGT 3'. The amplification products were resolved on 0.7% agarose-TAE gel for 1.5 h.

**Statistical Analysis**

The statistical analysis was performed in the GraphPad Prism 8 Software. Data are shown as media ± SEM, unless stated otherwise. Data normality was tested using Shapiro-Wilk's test. For multiple comparisons, normally distributed data were analyzed using a one-way ANOVA followed by Dunnett's multiple comparison test. For multiple nonparametric comparisons, a Kruskal-Wallis test followed by Dunn's multiple comparison test was used to determine significance. In all cases, data not indicated as significant should be considered not statistically different.

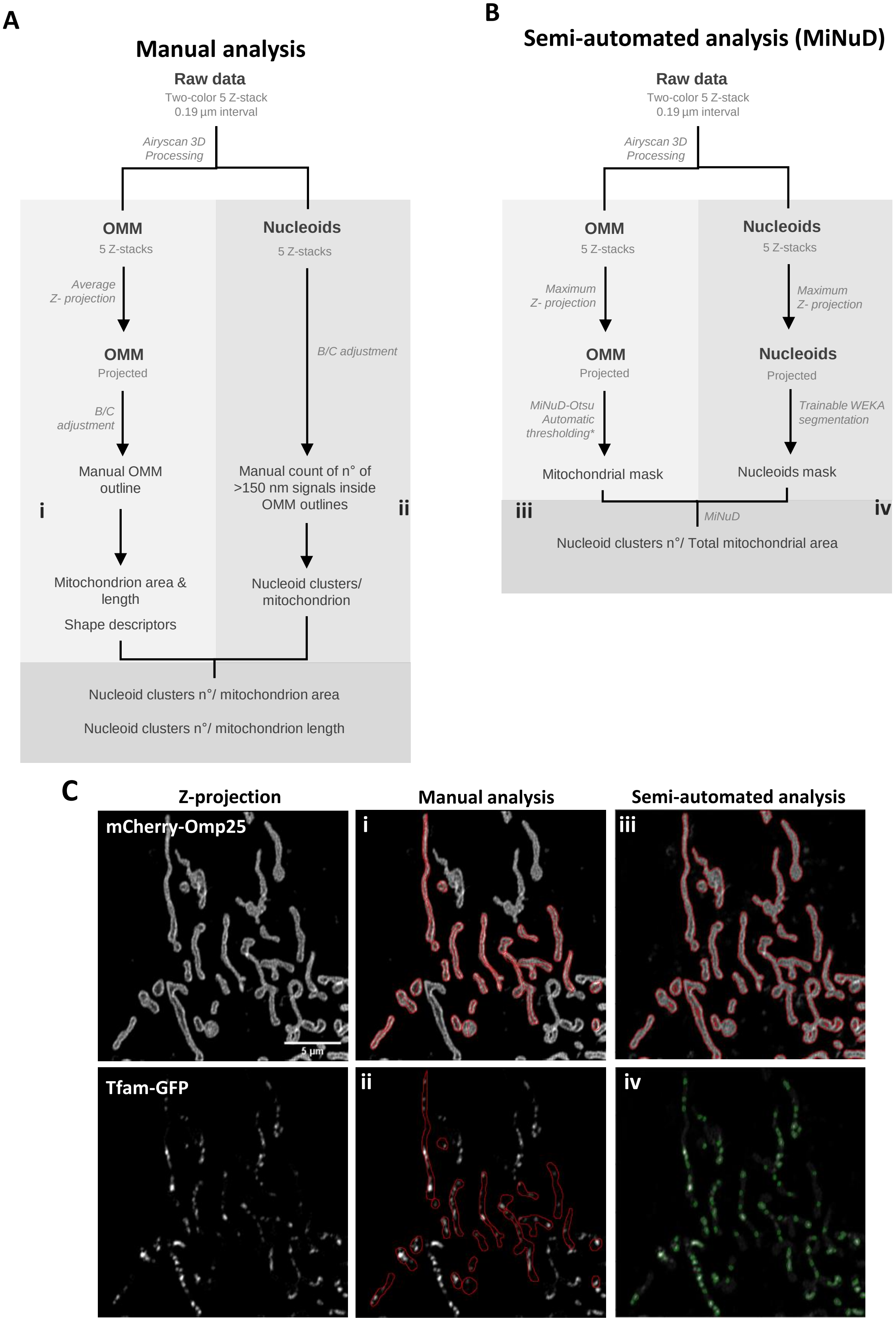


**Supplementary Figure 1. Manual and semi-automated (MiNuD) nucleoid cluster distribution analysis workflow. (A)** Schematic representation of the manual approach. 3D Airyscan processing was performed in ZEN Black software, and the following image processing steps were performed manually in FIJI ImageJ.

Mitochondrion signal eligibility criteria: no donut-shaped or other complex-shaped mitochondria were included; overlapping mitochondria were allowed if distinguishable as independent objects by 3D projection; the whole mitochondrial body should be inside the field of view. Nucleoids signal eligibility criteria: only nucleoid signals inside previously determined OMM outlines were quantified; the signals should be wider than 150 nm, considering the resolution limitations of our acquisition setting. The manual approach was applied to cells with mitochondria labeled with mCherry-Omp25 (OMM) and mitochondrial nucleoids labeled with Tfam-GFP. **(B)** Schematic representation of the semi-automated MiNuD approach. 3D Airyscan processing was performed in ZEN Black software, maximum Z projection and classification with Trainable Weka were performed in FIJI ImageJ. The steps performed by the Python MiNuD code in Anaconda are marked with an asterisk. The semi-automated approach was applied to cells with mitochondria labeled with mCherry-Omp25 or MitoTracker Deep Red, and nucleoids labeled with Tfam-GFP or PicoGreen. Each workflow is accompanied by a data processing example using a representative WT MEF cell co-transfected with mCherry-Omp25 and Tfam-GFP cDNA. (i) Red lines show the manual OMM outline (area) and inner line (length) of the mitochondria that fulfill the eligibility criteria. (ii) Red lines show the manual OMM outline used to discriminate which nucleoids to count and the mitochondrion they belong to. The information from (i) and (ii) was used to calculate nucleoid distribution parameters in Microsoft Excel. (iii) Red lines show the automated mitochondria masking performed by Otsu thresholding in the MiNuD code. (iv) Green lines show the semi-automated nucleoid masking performed by Trainable Weka in FIJI ImageJ. The masks shown in (iii) and (iv) were used as input for the distribution analysis in the MiNuD code.

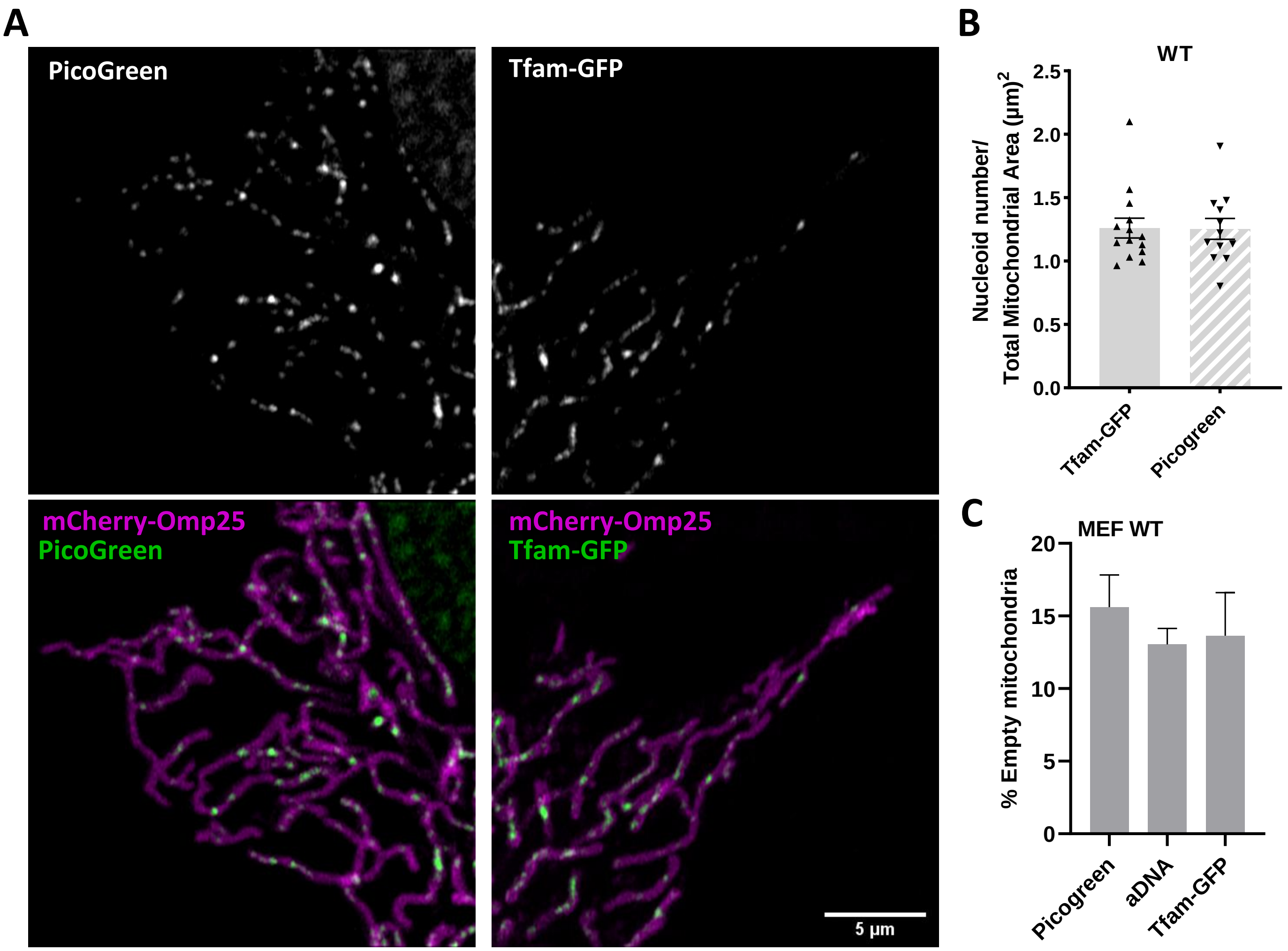


**Supplementary Figure 2. Tfam overexpression and PicoGreen staining display comparable nucleoid cluster distribution. (A)** Representative images of WT MEF transfected with mCherry-Omp25 (OMM) and labeled with PicoGreen or co-transfected with Tfam-GFP (Nucleoids). **(B)** Number of nucleoid clusters per total mitochondrial area, quantified semi-automatically in FIJI Image J/Python by the MiNuD algorithm. Data are mean ± SEM from ≥ 3 independent experiments. **(C)** Percentage of mitochondria without nucleoids or "empty" in live WT MEFs labeled with Picogreen or Tfam-GFP or fixed and labeled with anti-dsDNA. Data are mean ± SEM from ≥ 26 cells ≥ 3 independent experiments.

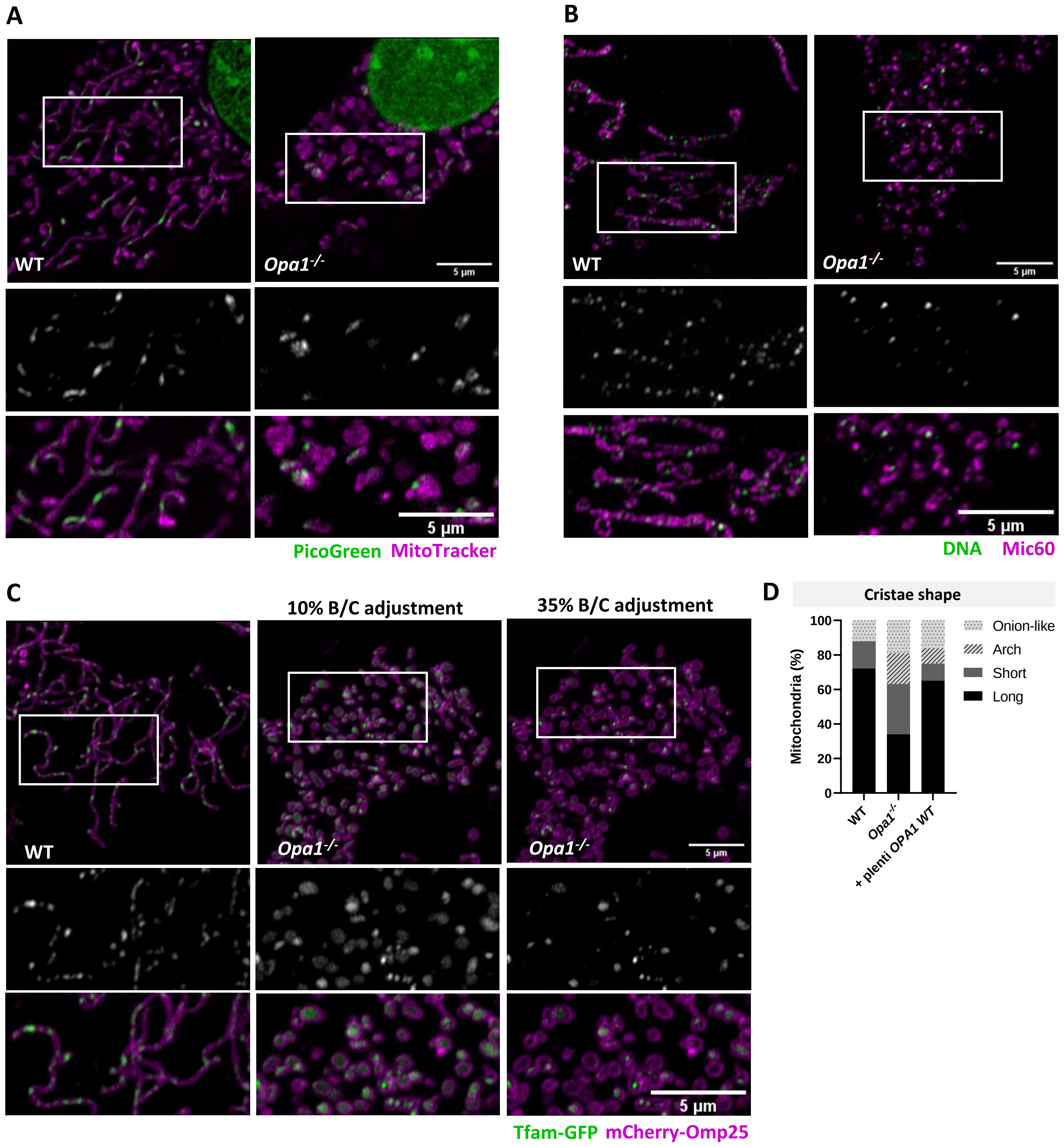


**Supplementary Figure 3. Diverse labeling strategies confirm nucleoid abundance reduction in *Opa1*$^{-/-}$ cells.** **(A)** Representative images of live-cell WT MEF and *Opa1*$^{-/-}$ labeled with MitoTracker Deep Red (Intermembrane space) and PicoGreen (DNA). **(B)** Representative images of fixed WT MEF and *Opa1*$^{-/-}$ labeled with anti-DNA and anti-Mic60 antibodies. **(C)** Representative images of live-cell WT MEF and *Opa1*$^{-/-}$, co-transfected with mCherry-Omp25 (OMM) and Tfam-GFP (nucleoids) cDNA. The central and right rows correspond to the same image with different Brightness/Contrast adjustments. The intense mitochondrial background noise of Tfam-GFP was filtered using a higher contrast adjustment of ~35% in FIJI ImageJ to obtain puncta resembling the nucleoid signals obtained through complementary labeling strategies in A-B. In each image, the bottom panel corresponds to an inset of the green channel (top) and merge (bottom). **(D)** Mitochondrial cristae characteristics classification in WT MEF, *Opa1*$^{-/-}$ and *Opa1*$^{-/-}$ cells stably expressing a lentiviral plasmid carrying OPA1 cDNA., separating aberrant cristae in onion-like and arch phenotypes. Data are from ≥ 98 objects from ≥ 2 independent experiments.

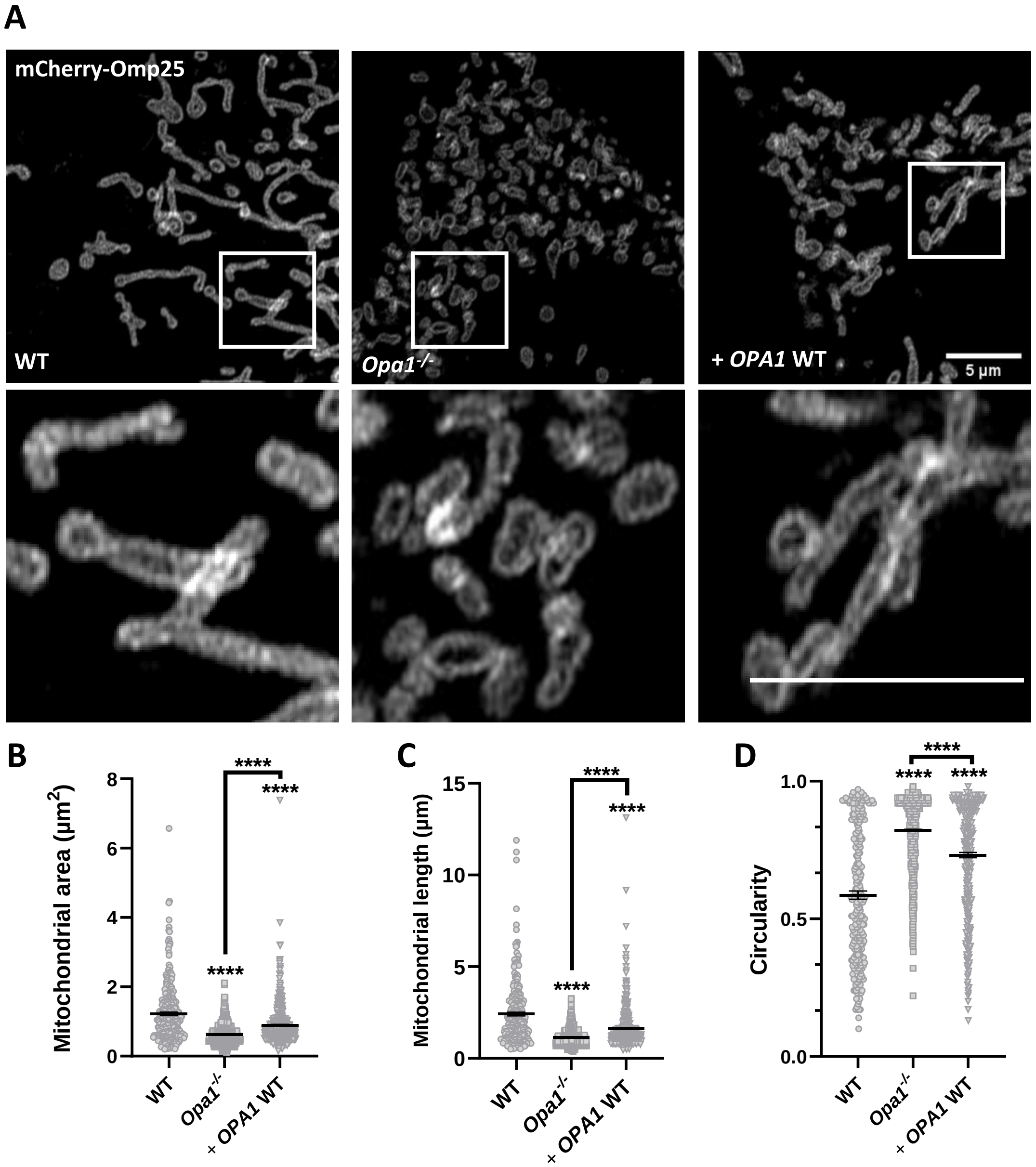


**Supplementary Figure 4. OPA1 acute expression partially rescues normal mitochondrial morphology**. **(A)** Representative images of MEF WT, *Opa1*$^{-/-}$ and *Opa1*$^{-/-}$ cells with WT OPA1 acute expression, co-transfected with mCherry-Omp25 (OMM) cDNA. Bottom panel, zoom-in of white squares. Mitochondrial morphology was assessed using **(B)** mitochondrial area, **(C)** mitochondrial length and **(D)** circularity. Data are mean of ≥ 281 objects from ≥ 15 cells of ≥ 3 independent experiments (**** $p<0.0001$).

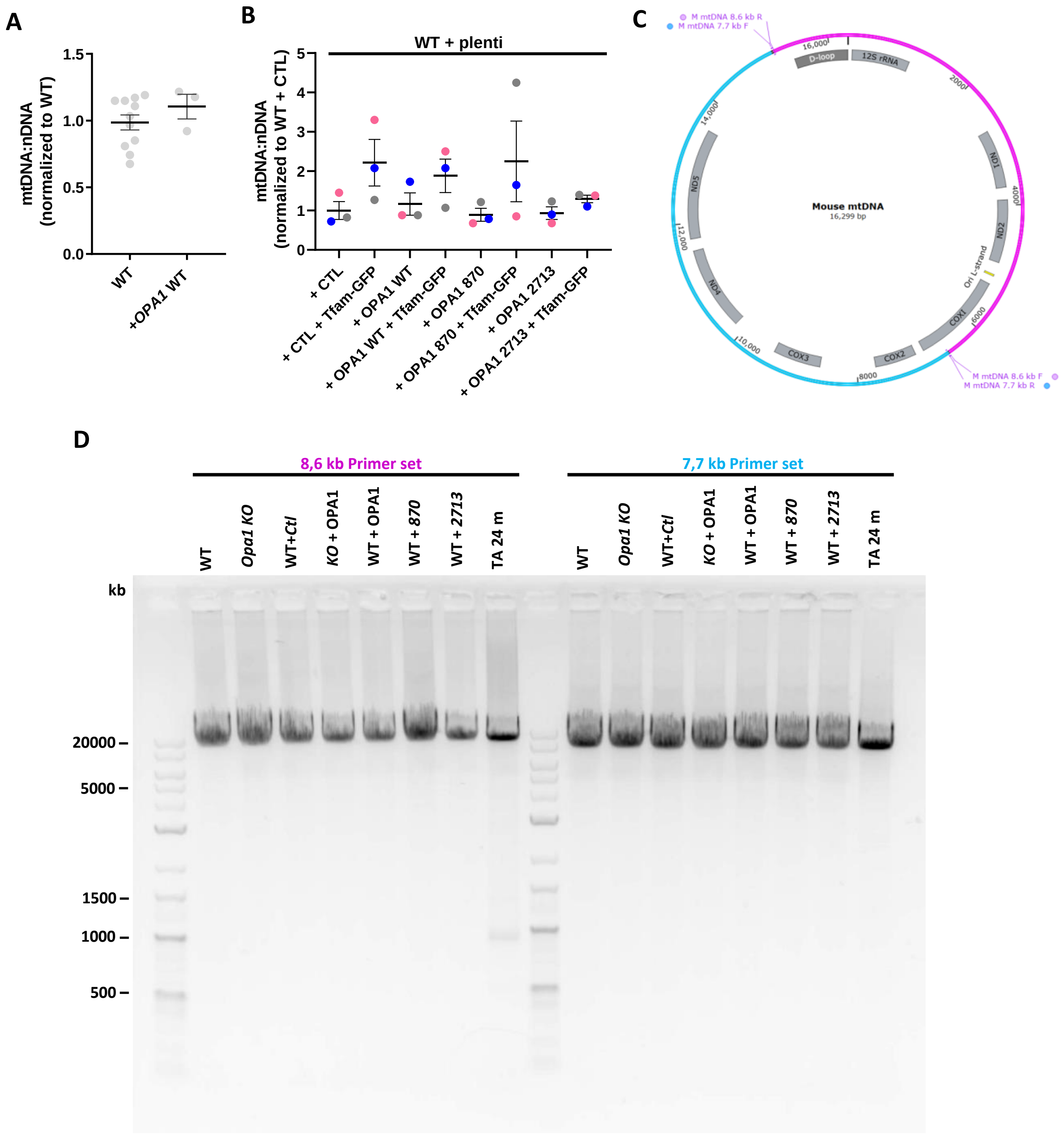


**Supplementary Figure 5. MtDNA levels are not significantly affected by OPA1 overexpression or Tfam-GFP, and cells stably expressing OPA1 variants don't show mtDNA deletions (A)** Mitochondrial DNA abundance of WT MEFs with the acute expression of *OPA1* WT quantified by qPCR of mt-Nd4 and Gadph ratio. Data are mean ± SEM from ≥ 3 independent experiments. **(B)** Mitochondrial DNA abundance of WT MEFs, *Opa1*$^{-/-}$ and WT MEFs with stable expression of OPA1 variants and acute expression of Tfam-GFP, quantified by qPCR of mt-Nd4 and Gadph ratio. Data are mean ± SEM from ≥ 3 independent experiments. **(C)** Mouse mtDNA map with the binding sites for primers to amplify an 8.6 kb (magenta) and 7.7 kb (cyan) fragments. **(D)** Representative Long-range PCR gel (2 independent experiments) of WT MEFs, *Opa1*$^{-/-}$ and WT MEFs with stable expression of OPA1 variants. A Tibialis Anterior extract from a 24-month-old mouse was added as a positive control for mtDNA deletions.

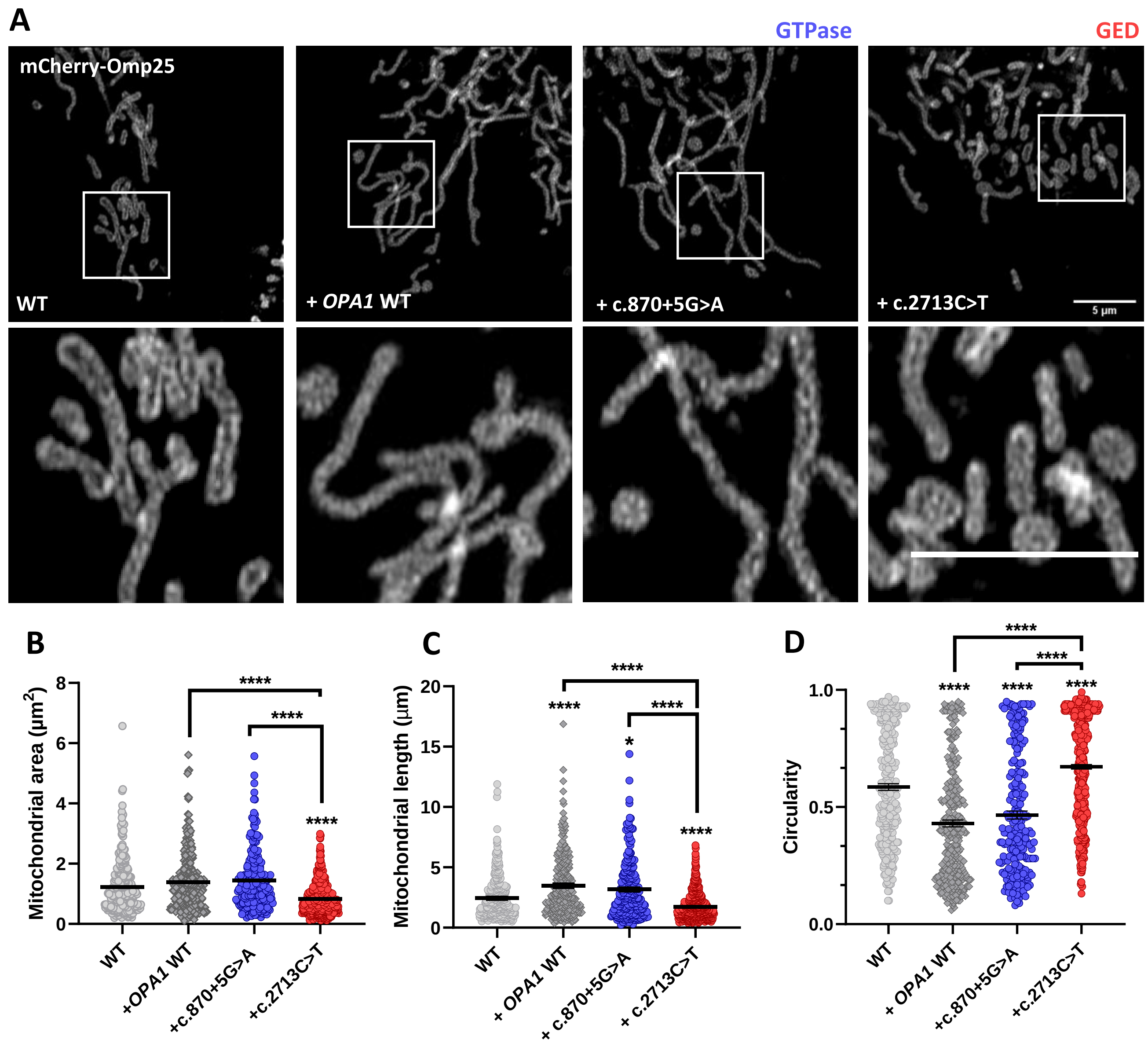


**Supplementary Figure 6. Acute expression of WT and pathogenic *OPA1* variants cause distinct mitochondrial morphology alterations. (A)** Representative images of MEF WT, MEF WT expressing OPA1 WT, and pathogenic OPA1 variants, co-transfected with mCherry-Omp25 (OMM) and Tfam-GFP (nucleoids) cDNA. Bottom panel, zoom-in of white squares. Mitochondrial morphology was assessed using **(B)** mitochondrial area, **(C)** mitochondrial length, and **(D)** circularity. Data are mean ± SEM of ≥ 228 objects from ≥ 18 cells of ≥ 4 independent experiments (**** $p<0.0001$; * $p<0.05$).
